# Uncovering topologically associating domains from three-dimensional genome maps with TADGATE

**DOI:** 10.1101/2024.06.12.598668

**Authors:** Dachang Dang, Shao-Wu Zhang, Kangning Dong, Ran Duan, Shihua Zhang

## Abstract

Topologically associating domains (TADs) emerge as indispensable units in three-dimensional (3D) genome organization, playing a critical role in gene regulation. However, accurately identifying TADs from sparse chromatin contact maps and exploring the structural and functional elements within TADs remain challenging. To this end, we develop a graph attention auto-encoder, TADGATE, to accurately identify TADs even from ultra-sparse contact maps and generate the imputed maps while preserving or enhancing the underlying topological structures. TADGATE can capture specific attention patterns, pointing to two types of units with different characteristics in TADs. Moreover, we find that the organization of TADs is closely associated with chromatin compartmentalization, and TAD boundaries in different compartmental environments exhibit distinct biological properties. We also utilize a two-layer Hidden Markov Model to functionally annotate the TADs and their internal regions, revealing the overall properties of TADs and the distribution of the structural and functional elements within TADs. At last, we apply TADGATE to highly sparse and noisy Hi-C contact maps from 21 human tissues or cell lines, enhancing the clarity of TAD structures, investigating the nature of conserved and cell type-specific boundaries, and unveiling the cell type-specific transcriptional regulatory mechanisms associated with topological domains.

## Introduction

The continuous improvement of chromosome conformation capture technologies and their derivatives, such as Hi-C, has facilitated the generation of a significant volume of 3D genomics data (1–5). These datasets span multiple species and cell types (6–9), encompassing various biological processes, such as neural development (8) and somatic cell reprogramming (10), thus providing unprecedented opportunities to unravel the mysteries of 3D genomic organization within the cell nucleus. The genome adopts a hierarchical arrangement with multiscale 3D chromatin structures, including chromosomal territories (4, 11), A/B compartments and subcompartments (4, 5), topologically associating domains (TADs) (12, 13), and chromatin loops (5).

TADs appear as squares with increased intensity along the diagonal of a Hi-C contact map, representing structural domains with enhanced self-interactions. TAD boundaries, the start and end regions of a domain, serve as insulators to prevent inter-TAD interactions and favor intra-TAD interactions (13). The investigation of boundaries has been a focal point in TAD research (14–16). The boundaries are enriched with chromatin insulator protein CTCF, the cohesin complex (RAD21 and SMC3), and housekeeping genes (12). Moreover, the positions of TAD boundaries are highly conserved across various cell types and even among different species (5–7, 12, 14, 17). TADs are often considered stable neighborhoods for gene regulation (18), and the disruptions in TAD boundaries caused by genomic structural variations may lead to severe diseases (19, 20), including cancer (21, 22). Therefore, TADs are the basic structural and functional units of chromatin organization, and the accurate determination of TADs is critical for 3D genomic research.

Researchers have developed computational methods to identify TADs based on diverse strategies. The most classical approaches involved calculating one-dimensional (1D) indicators from chromatin contact maps, such as Directionality Index (DI) (12) and Insulation Score (IS) (13), and then identifying TAD boundaries based on the specific indicator patterns. Later, TopDom (23) and OnTAD (24) were further designed based on modified 1D indicators. Moreover, MSTD (25) and IC-Finder (26) extracted the interaction signals from the Hi-C contact map and adopted specific clustering methods to obtain TADs. Besides, some methods, including 3DNetMod (27) and deDoc (28), treated the Hi-C contact map as an adjacency matrix of chromatin interaction network and utilized community detection or graph segmentation algorithms for this task. Despite employing diverse computational frameworks, these methods still face challenges in accurately identifying TADs from sparse and noisy contact maps of the Hi-C dataset with insufficient sequencing.

Zufferey et al. (29) evaluated various methods using down-sampled Hi-C datasets to simulate inadequate sequencing depth. Apart from IS and TopDom, many methods struggled to identify TADs robustly in sparse contact maps. Lee and Roy developed GRiNCH (30) to simultaneously detect topological domains and smooth the sparse chromatin contact matrices with graph-regularized non-negative matrix factorization. However, they merely down-sampled the Hi-C dataset from the cell line with higher sequencing depth to levels of other cell lines with slightly lower depth (30). They investigated the robustness of GRiNCH in identifying TADs at different sequencing depths without fully considering its performance on sparse contact matrices when the datasets suffer from insufficient sequencing. They didn’t intuitively display or evaluate how this method contributed to the smoothness or imputation of the sparse contact matrices. However, since the advent of Hi-C, due to the limitations of this technology and the poor quality of biological samples, datasets with relatively lower sequencing depth have been generated and accumulated (6, 10). These datasets could generate sparse and noisy contact matrices, hindering subsequent structure identification and biological analyses. Hence, it is essential to develop computational methods capable of imputing and smoothing the sparse Hi-C contact matrices while accurately identifying structural units like TADs. That is crucial for fully leveraging the potential of these valuable datasets.

Currently, most studies on TADs focus on their boundaries. However, they often treat all boundaries equally and analyze their collective properties, such as CTCF enrichment, without distinguishing between them (12, 23). More and more evidence indicates that TAD boundaries are categorized into different types, each exhibiting unique characteristics (16, 24, 31). Therefore, exploring the properties of different TAD boundaries may deepen our understanding of the organizational principles of TADs. More importantly, TAD boundaries only occupy a small portion of the genome, and many genes and regulatory elements are located within the domain interior. However, TAD still functions like a black box. The internal structural or functional elements and their mutual interactions remain unclear. Jiang et al. (32) defined the spatial density of open chromatin (SDOC) for each TAD by combining its 3D volume with the total number of accessible chromatin regions. They demonstrated that TADs with higher SDOC were more likely to be active domains. Besides, our previous study indicated that TADs could be divided into diverse types based on the DNA replication timing, where TADs replicating earlier tended to be active domains with high-level gene expression (31). These findings indicate the importance of more detailed classification and research on both TAD boundaries and the entire domains, and the structural and functional elements within TADs also deserve further exploration.

To this end, we present TADGATE to identify <u>TAD</u>s and perform smoothing and imputation of Hi-C contact maps with a <u>G</u>raph <u>AT</u>tention auto-<u>E</u>ncoder. TADGATE substantially improves the accuracy of TAD identification and denoises the contact maps while preserving or enhancing the topological domains. The model learns two distinct patterns, i.e., the attention valleys provide potential sites for TAD boundaries, and the attention peaks correspond to functional elements within the domains. These patterns exhibit different histone modifications and chromatin states. We also elucidate the close relationship between TADs and chromatin compartmentalization and categorize TAD boundaries into three types based on their compartmental environments. We demonstrate that different types of boundaries differ in the insulation strength for chromatin contact, the content of repetitive DNA elements, gene expression level, and the distribution of epigenomic markers. We annotate TADs and their internal structural or functional elements using a two-layer hidden Markov model combined with multiple epigenomic markers. Finally, leveraging TADGATE, we obtain clearer TADs in sparse chromatin contact maps from 21 human tissues and cell lines and depict the conserved and cell type-specific TAD boundaries relating to cell type-specific gene transcription regulation.

## Results

### Overview of TADGATE

Each genomic bin in the input Hi-C contact map is treated as a sample, and its interaction vector with other bins serves as the sample feature. Considering that a TAD consists of several continuous genomic bins, TADGATE first constructs a neighborhood graph to represent the proximity relationships between bins along the 1D genome based on a pre-defined radius (**Fig. 1**). In the neighborhood graph, each genomic bin is connected only to upstream or downstream bins within the radius. TADGATE learns a low-dimensional latent embedding for each genomic bin with genomic proximity and chromatin interactions via a graph attention auto-encoder. More specifically, the interaction vector of each bin is first transformed into a latent embedding by an encoder and then reversed back into a reconstructed interaction profile via a decoder. All these reconstructed interaction profiles can form the imputed contact map, and the latent embeddings are used to visualize these bins with UMAP (33) (**Fig. 1**).

**Figure 1.**
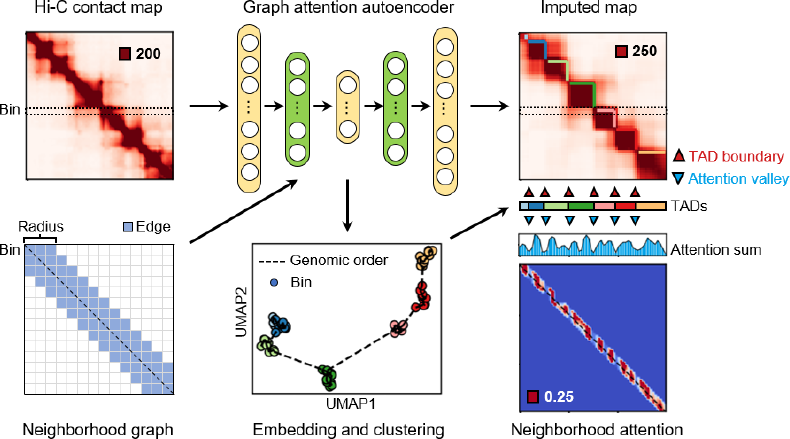
Overview of TADGATE. TADGATE employs a graph attention auto-encoder to detect topologically associating domains (TADs) from Hi-C contact maps and reconstructs refined maps that preserve or enhance the chromatin domain structure. TADGATE first constructs a neighborhood graph to represent the proximity relationships between genomic bins and further learns low-dimensional latent representations by harnessing genomic proximity and chromatin interactions via a graph attention auto-encoder. The input of the auto-encoder is the chromatin interaction vector of each bin, and the graph attention layer is adopted in the middle of the encoder and decoder (green layers). The model output consists of the imputed map, the embedding vector for each bin, and the attention map. The imputed map offers a smoothed version of the original Hi-C contact map that retains the topological structures. The embedding vectors facilitate clustering bins into discernible TADs. The high degree of concordance between TAD boundaries and attention valleys in the attention sum profile also demonstrates the model’s capacity to capture and depict the topological information within the Hi-C contact map.

By adopting an attention mechanism, TADGATE adaptively learns the attention weights of each edge in the neighborhood graph and uses them to update the bin representation by collectively aggregating information from neighbor bins. Thus, we get the neighborhood attention map, in which the attention weights reflect the similarity of topological information between neighbor bins. When summing up the attention weights by columns in the neighborhood attention map, we observe that the attention valleys match well with the TAD boundaries in the Hi-C contact map (**Fig. 1**), indicating the attention mechanism has learned valuable information in identifying topological domains. Finally, we combine the input and imputed Hi-C maps, the embeddings of bins, and the patterns in the attention sum profile to identify topological domains (**Methods and Supplementary Fig. S1**).

### TADGATE identifies more TADs with high confidence

We compared TADGATE with four competing methods using the Hi-C data from the GM12878 cell line. These methods included two classic ones, DI and IS, and TopDom, which consistently ranked high in many benchmarking studies (29, 30). We also included GRiNCH, which could provide a smoothed contact map while identifying TADs. We found that TADGATE could identify more TADs than other methods, and the average domain length was around 300kb (**Fig. 2a and Supplementary Fig. S2a**), suggesting that TADGATE can identify a finer-scale organization of the topological domains. Each TAD can serve as a cluster of bins with similar chromatin contact patterns. The silhouette coefficient and Davies-Bouldin index showed that TADGATE could achieve more reasonable cluster divisions of bins, implying better TAD identification than other methods (**Fig. 2B and Supplementary Fig. S2B**). Besides, we found a clear enrichment of CTCF binding at the boundaries of all methods. The enrichment was most pronounced at the boundaries of DI and TopDom, followed by TADGATE, which was more significant than IS and GRiNCH (**Fig. 2C**). We also observed similar patterns for the two components of cohesin, RAD21, and SMC3 (**Supplementary Fig. S2C**).

**Figure 2.**
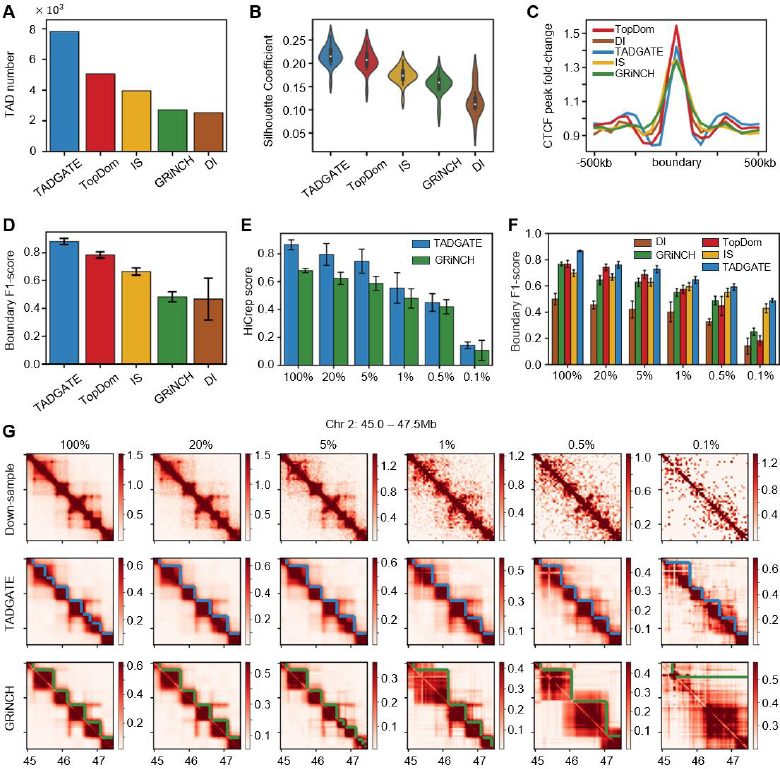
Comparison of TAD-calling methods. (**A**) Number of TADs identified by different methods in chromosome 1-22 and X of GM12878 cell line. (**B**) Silhouette coefficients of TADs by various methods on each chromosome. (**C**) CTCF binding profiles around the TAD boundaries of different methods. (**D**) F1-score of boundaries identified by diverse methods on each chromosome. (**E**) HiCrep scores between TADGATE-imputed or GRiNCH-imputed maps and experimental Hi-C contact maps (with 100% reads) at different down-sampling ratios. (**F**) F1-score of boundaries identified by various methods at different down-sampling ratios. (**G**) Hi-C contact maps of a region on chromosome 2 at various down-sampling ratios. The corresponding maps imputed by TADGATE and GRiNCH, and the TADs identified by them are shown. The TADs identified by other methods are shown in Supplementary Fig. S3F.

We further performed pairwise comparisons between TADGATE and each of the other four methods (**Supplementary Fig. S2F-I**). For each of the four methods, over 80% of the boundaries they identified could be captured by TADGATE. Conversely, many TADGATE boundaries were challenging to detect by competing methods, especially for DI and GRiNCH. Moreover, we found that the unique boundaries identified by TADGATE showed clear CTCF enrichment and insulation effects on chromatin contacts. However, the unique boundaries of DI, IS, and GRiNCH showed abnormal patterns, indicating that these boundaries might be false positive ones or the positions were not accurate (**Supplementary** Fig. S2F, G, I). As for TADGATE and TopDom, TADGATE missed some boundaries unique to TopDom, but that accounted for a tiny portion. TADGATE could capture more boundaries with slightly weaker CTCF enrichment and localized insulation effects than TopDom, indicating a great sensitivity to boundary detection of TADGATE (**Supplementary Fig. S1H**). That also explained why the average enrichment of CTCF at the boundaries of TADGATE was weaker than TopDom (**Fig. 2C**).

In addition to the pairwise comparison, we constructed a unified boundary set according to the boundary voting strategy used in our previous study (31) (Methods and **Supplementary Fig. S1C**). The unified boundaries in this set represent bins identified as TAD boundaries by at least one method and exhibit enrichment of CTCF and insulation effects on chromatin contacts. Thus, they can serve as highly reliable boundaries for comparing the precision and recall of different methods. TADGATE showed similar boundary precision to other methods and significantly higher boundary recall (**Supplementary Fig. S1D, E**). Overall, TADGATE exhibited a notably higher boundary F1-score than other methods, indicating its ability to comprehensively identify TADs from Hi-C contact maps while maintaining high accuracy (**Fig. 2D**).

### TADGATE simultaneously imputes sparse contact maps and robustly identifies TADs

We applied TADGATE and four competing methods to the down-sampled Hi-C contact maps of the K562 cell line with sampling ratios of 20%, 5%, 1%, 0.5%, 0.1% (**Fig. 2G** and **Supplementary Fig. S3A**). TADGATE and GRiNCH could generate the imputed Hi-C contact maps. The sparsity of the TADGATE-imputed maps was significantly reduced compared to the down-sampled ones (**Supplementary Fig. S3A**). GRiNCH assigns a value to each pixel of the contact map, and the sparsity of all the imputed maps is zero. We used the HiCrep score (34), peak signal-to-noise ratio (PSNR), and structural similarity (SSIM) (35) to evaluate the similarity between the imputed map and the contact map with 100% reads. The HiCrep scores became smaller as the down-sampling ratio decreased for TADGATE and GRNiCH. But the contact maps imputed by TADGATE consistently exhibited higher HiCrep scores (**Fig. 2E**). At 100% sampling ratio, we observed higher values of the PSNR and SSIM for TADGATE, implying that TADGATE outperformed GRiNCH in preserving topological information during the map imputation. Besides, at the other down-sampling ratios, TADGATE also achieved robust and high values of PSNR and SSIM, indicating its capability to enhance or reconstruct the topological structures in sparse contact maps (**Supplementary Fig. S3B**). We also used the Jaccard index to assess the overlap of TAD boundaries between down-sampled maps and the full-reads map. We observed decreased boundary overlap for all methods as the sampling ratio decreased. TopDom, IS, and TADGATE performed relatively well with sampling ratios of 20% and 5%. TADGATE showed better robustness and could maintain a certain degree of boundary overlap in extremely sparse contact maps with 0.1% reads as the sampling ratio continued to decrease (**Supplementary Fig. S3C**). Subsequently, we utilized the TAD boundaries identified by the five methods in the contact maps with 100% reads to construct a unified boundary set and evaluated the boundaries of each method across different down-sampling ratios (**Fig. 2F and Supplementary Fig. S3D, E**). TADGATE achieved significantly higher boundary F1 scores across various down-sampling ratios than other methods, indicating its ability to identify TADs in sparse contact maps. We gave example regions to show the contact maps imputed by TADGATE and GRiNCH and the TADs identified by each method at different down-sampling ratios (**Fig. 2G, Supplementary Figs. S3F and S4**). TADGATE effectively imputed and smoothed the contact maps while preserving the internal topological structures (**Fig. 2G**). It robustly and accurately identified TADs across various down-sampling ratios and even enhanced and partially reconstructed the latent topological domains at a sampling ratio of 0.1%, where structures in the contact map were barely discernible. GRiNCH maintained good map imputations at higher down-sampling ratios, but its performance progressively deteriorated with reduced sampling. The TADs identified by GRiNCH were also adversely affected (**Fig. 2G**). TopDom and IS performed well in identifying TADs across multiple down-sampling ratios, but they became unstable when the contact maps were exceptionally sparse. DI tended to capture obvious TADs but might miss more underlying boundaries, and it failed on sparse contact maps (**Supplementary Fig. S3F and S4**).

In addition, we defined the boundary strength to reflect the insulating effect of boundaries on the interactions between chromatin on either side (**Supplementary Fig. S5A**). Next, we examined the boundary strength of the previously defined unified boundary set in various down-sampled and imputed maps. Except in very sparse contact maps, the boundary strengths of these unified boundaries were enhanced in both the TADGATE-imputed and GRiNCH-imputed maps when compared with the down-sampled maps (**Supplementary Fig. S5B, C**). It suggested that both methods could strengthen the topological domains, making them more detectable. All these results indicated that TADGATE could generate the imputed or smoothed contact maps with clear structures and identify TADs accurately and robustly.

### Attention peaks and valleys refer to units with distinct properties within TADs

The attention map in TADGATE exhibits specific structures. When summing up the values in the attention map by column, we observed a strong correspondence between the attention valleys and the TAD boundaries in the Hi-C contact map (**Fig. 1 and Supplementary Fig. S6A**). The bins belonging to the same TAD also had similar latent embeddings learned by TADGATE (**Fig. 3A** and **Supplementary Fig. S6B, C**). We collected all valleys and peaks in the attention sum profile and obtained the aggregated Hi-C contact maps around them. The attention valleys corresponded to the domain boundaries, and the attention peaks were located within the domains (**Fig. 3B**). The attention peaks were primarily located near the middle of domains but not domain centers (**Supplementary Fig. S6D**). The attention peaks exhibited stronger chromatin interactions with other bins within the same domain, and the interactions between attention valleys and other bins were much weaker (**Fig. 3C**). However, genes residing in attention valleys exhibited significantly higher expression levels than genes in attention peaks (**Fig. 3D**).

**Figure 3.**
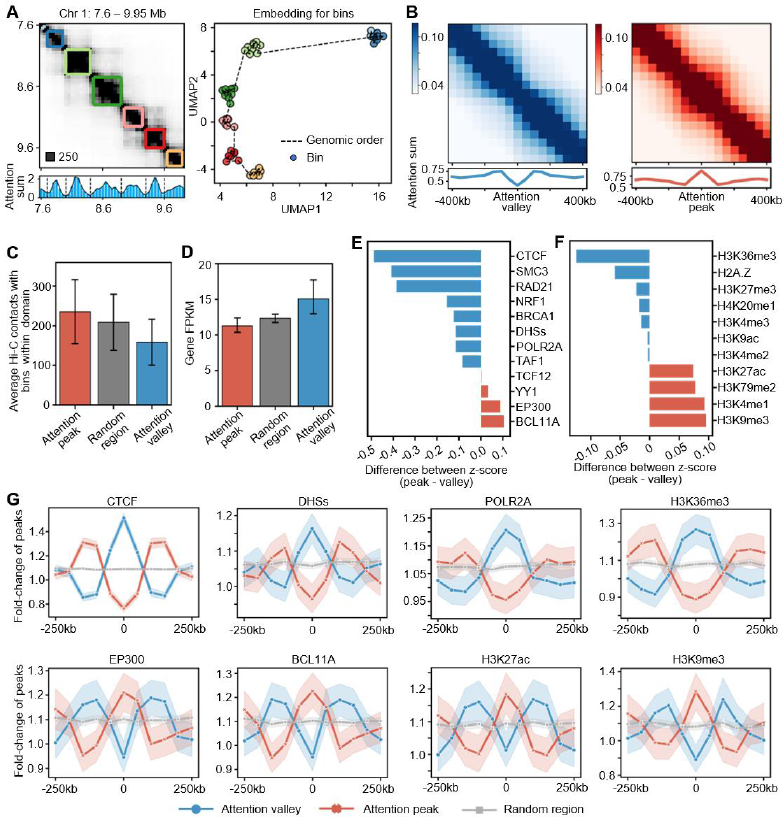
Attention peaks and valleys facilitate the discovery of two types of distinct structural and functional elements within TADs. (**A**) Hi-C contact map of a region on chromosome 1 of GM12878, along with the TADs identified by TADGATE. The corresponding profile of attention sum is shown below, and dashed lines mark the attention valleys. The UMAP plot on the right is generated based on the embedding vectors of bins obtained from TADGATE, with bins colored according to the TADs they belong to. (**B**) Aggregated Hi-C contact maps around attention valleys and peaks and the averaged attention sum profiles. (**C**) Average Hi-C contacts between attention valleys or peaks and other bins within the same domain. Random regions, excluding attention valleys and peaks, were selected as controls. (**D**) Expression patterns of genes located at attention valleys, attention peaks, and randomly selected regions. (**E**) The difference in enrichment z-scores of several transcription factors between attention peaks and valleys. (**F**) The difference in enrichment z-scores of several histone modifications between attention peaks and valleys. (**G**) Profiles of some transcription factors, histone modifications, and DHSs around attention peaks or valleys. The shaded area represents the 95% confidence interval in bootstrap.

We also observed significant differences in the distribution of various transcription factors (TF) and histone modifications between attention peaks and valleys. CTCF, cohesin components RAD21 and SMC3, and H3K36me3 and H3K4me3, which serve as markers of active gene expression, exhibited significantly stronger signals on attention valleys. Meanwhile, attention peaks showed stronger signals for EP300, BCL11A, H3K27ac, and H3K9me3 (**Fig. 3E, F**). We also identified the enriched TF motifs in the accessible regions of attention peaks and valleys, with the most notable difference being a higher enrichment of CTCF and CTCFL in attention valleys (**Supplementary Fig. S6E, F**). We found that some epigenetic modifications exhibited opposite distribution patterns around the two types of units (**Fig. 3G and Supplementary Fig. S7A**). For example, the CTCF, DNase I Hypersensitive Sites (DHSs), POLR2A, and H3K36me3 were significantly enriched near attention valleys but showed a marked reduction in attention peaks. EP300, BCL11A, H3K27ac, and H3K9me3 exhibited a contrasting pattern. They significantly enriched around attention peaks but depleted near attention valleys (**Fig. 3G**). We also explored the distribution of 15 chromatin states annotated by ChromHMM (36) in attention peaks and valleys. The attention valleys were enriched in chromatin states related to active gene expression, including Strong transcription, Active TSS, etc. In contrast, attention peaks were enriched in chromatin states with regulatory functions such as heterochromatin and enhancers (**Supplementary Fig. S7B**). In summary, attention valleys serve as structural units to bound topological domains, containing some active genes with fewer associations with other regions within the domain. On the other hand, attention peaks act as core units interacting with other members and function as regulatory elements, which can affect the remaining regions in the same domain.

### TAD organization relates to chromatin compartmentalization

Chromatin compartments and TADs depict different levels of chromatin organization, representing global chromatin partitioning and local structural units, respectively. However, they still exhibit certain relationships in some studies (37, 38). Herein, based on a more refined characterization of topological domains by TADGATE, we found a large proportion of TADs showed high purity in compartment composition, with over 80% entirely belonging to either compartment A or B (**Fig. 4A**). When considering finer chromatin subcompartments, 67.4% of TADs only contain one type of subcompartment and very few TADs (∼3%) span more than two different subcompartments (**Fig. 4B**). Moreover, the switch points between two different types of compartments or sub-compartments show substantial overlap with TAD boundaries. Nearly 40% of compartment switch points and 35% of subcompartment switch points coincide with TAD boundaries (**Fig. 4C**). Overall, the distance between compartment or subcompartment switch points and TAD boundaries is significantly smaller than those for randomly shuffled boundaries, implying a close relationship between these two different scales of chromatin structures (**Supplementary Fig. S8A, B**).

**Figure 4.**
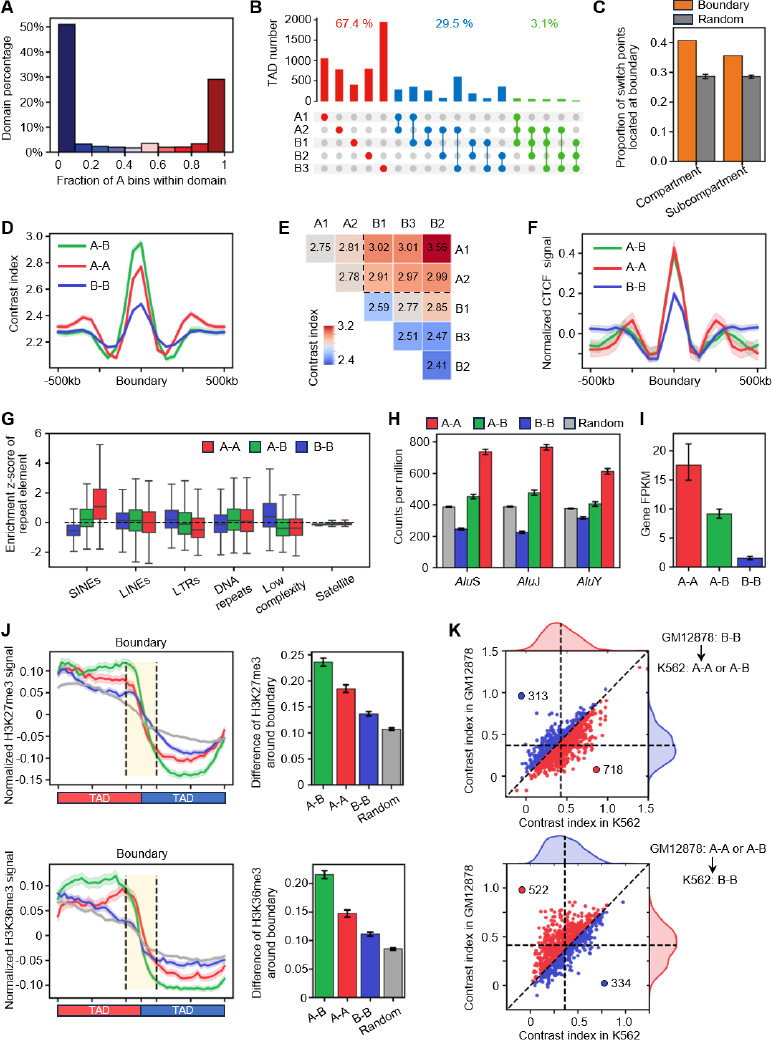
TAD organization is correlated with chromatin compartmentalization. (**A**) Distribution of domains with various fractions of bins belonging to compartment A. (**B**) The number of TADs with distinct subcompartment compositions. TAD comprises a subcompartment if it occupies at least 10% of the TAD region. (**C**) Proportion of compartment or subcompartment switch points located at TAD boundaries. We performed 100 random permutations of TADs as controls. (**D**) Comparison of the contrast index of TAD boundaries between different combinations of compartments. (**E**) Average contrast index of TAD boundaries between different combinations of subcompartments. The region marked by dashed lines in the upper right corner indicates the results between subcompartments belonging to types A and B. (**F**) The normalized CTCF signal around TAD boundaries between different combinations of compartments. (**G**) The enrichment z-score of repeat elements at TAD boundaries between different combinations of compartments. (**H**) Density of the *Alu* subfamilies in TAD boundaries between different combinations of compartments. (**I**) Expression patterns of genes located at TAD boundaries between different combinations of compartments. (**J**) The profile of H3K27me3 and H3K36me3 around TAD boundaries between different combinations of compartments. For each boundary, the direction of the profile is adjusted to keep a higher signal in the left TAD than the right one. The bar plot on the right side exhibits the difference between the average signals within the two TADs. (**K**) The distribution of contrast index of TAD boundaries exhibiting conserved positions but different compartment types between GM12878 and K562. The boundaries with higher contrast index in each cell line are marked with red or blue and the corresponding boundary numbers are also shown. The vertical and horizontal dashed lines indicate the mean contrast index of these boundaries in GM12878 and K562.

Furthermore, TAD boundaries in different compartmental environments exhibit distinct insulation effects on chromatin contacts (**Fig. 4D**). The TAD boundaries that coincide with the switch points between the A and B compartments exhibit stronger insulation effects than boundaries between TADs within the same type of compartments. The boundaries between active TADs in compartment A are stronger than those between repressive TADs in compartment B. That may be attributed to the involvement of more gene regulatory events in active TADs, where stronger boundaries can protect the regulatory processes from external influence. The same patterns hold for TAD boundaries across two subcompartments. The TAD boundaries bridging A-type and B-type subcompartments provide the strongest insulation against chromatin interactions, and boundaries between A-type subcompartments typically demonstrate greater insulating properties than those between B-type subcompartments. (**Fig. 4E and Supplementary Fig. S8D**). The A1 type represents the most active subcompartment, while B3 and B2 are repressive subcompartments. A2 and B1 act as transitional subcompartments between them (39). The TAD boundaries became progressively stronger as the flanking subcompartment types showed higher discrepancies (**Supplementary Fig. S8E**).

Following this, we categorized the TAD boundaries into three types based on their compartmental environments, A-B, A-A, and B-B, and further explored their biological relevance. First, we identified the top 20 TF motifs enriched in the accessible regions of the three types of boundaries using Homer (40). We observed a notable similarity among the enriched motifs across the three types of boundaries, with CTCF and CTCFL consistently standing out as the most enriched motifs (**Supplementary Fig. S8F)**. When expanding the analysis to the top 100 enriched motifs, we still observed a high degree of overlap in enriched motifs among the three types of boundaries (**Supplementary Fig. S8G)**. Despite the similarity in enriched motifs, different types of boundaries exhibit varying degrees of chromatin accessibility. The A-A and A-B-type boundaries are notably more accessible than the B-B-type ones. Correspondingly, CTCF, RAD21, and SMC3 also show more significant enrichment at A-A and A-B-type boundaries than others (**Fig. 4F** and **Supplementary Fig. S8H)**. Besides, the distribution of repetitive DNA sequences among diverse types of TAD boundaries is also different. The A-A-type boundaries are significantly enriched in short interspersed nuclear elements (SINEs), and B-B-type boundaries tend to have more long terminal repeats (LTRs) and low complexity repeats (**Fig. 4G**). *Alu*, the most prevalent SINE, is widely distributed in the human genome and is enriched at TAD boundaries (7, 12, 31). We further observed that the A-A-type boundaries exhibited strong enrichment for three subfamilies of *Alu*, the A-B-type boundaries only showed weak enrichment, and the B-B-type boundaries lacked all subfamilies of *Alu* (**Fig. 4H**).

The expression levels of genes in A-A-type boundaries are significantly higher than those in A-B or B-B-type boundaries, and the housekeeping genes are also predominantly enriched in A-A-type boundaries (**Fig. 4I** and **Supplementary Fig. S8I**). Rao et al. reported that the distribution of histone modifications correlated with TAD structure (5). Additionally, we observed similar associations for various transcription factors. The signals of diverse histone modifications and transcription factors between regions within the same TAD exhibit stronger correlations, while the correlations between regions belonging to different TADs are significantly weakened. However, the RNA-seq signal is not highly correlated within domains, suggesting the presence of more intricate regulatory mechanisms for gene expression within TADs (**Supplementary Fig. S9A**). TADs show a stronger packaging effect on histone modifications than transcription factors (**Supplementary Fig. S9B**). We observed substantial divergence in histone modification and transcription factor signals flanking the A-B-type boundaries, followed by the A-A-type boundaries. There were also slight differences in epigenomic signals around the B-B-type boundaries (**Fig. 4J and Supplementary Fig. S9C**). It suggests that the organization of TADs and the strength of boundaries relate to surrounding epigenomic modifications and chromatin states.

Then we compared the TAD boundaries between GM12878 and K562. Almost 80% of boundaries remain conserved in position, and the majority of these conserved boundaries exhibit similar compartment types between GM12878 and K562 (**Supplementary Fig. S10B**). These compartment type-stable boundaries indicate no significant preference trend in contrast index between the two cell lines (**Supplementary Fig. S10C, D**). However, we also observed some boundaries undergoing compartment-type transitions between GM12878 and K562. Some boundaries are classified as B-B type in GM12878, but transiting to A-A or A-B types in K562. A greater proportion of these boundaries is associated with an increase in boundary strength compared to the compartment-type stable boundaries (Fisher’s exact test, *p*-value=1.44×10^-33^). In contrast, a higher proportion of boundaries shifting from A-A or A-B type in GM12878 to B-B type in K562 exhibit a lower contrast index (Fisher’s exact test, *p*-value=1.35×10^-9^) (**Fig. 4K and Supplementary Fig. S10E**). It suggests that the alterations in chromatin compartmentalization among different cell types are indeed related to changes in TAD organization and boundary strength.

### Functional annotation of TADs and internal regions with epigenomic markers

We employed a two-layer Hidden Markov Model (HMM) (41) to annotate TADs and their internal regions with multiple epigenomic markers in GM12878 (**Methods**). We defined 20 types of structural and functional elements within TADs, such as Promoter/enhancer, Structural element, and H3K36me3-dominant region, each exhibiting a distinct epigenetic modification pattern (**Fig. 5A and Supplementary Fig. S11B**). We discovered distinct distribution patterns of elements within TADs. For instance, some elements, such as Structural promoter/enhancer-2, Structural element, and H3K36me3-dominant region, tended to be located at the boundaries of TADs. In contrast, elements like Promoter/enhancer-1, Putative enhancer-1, and Transcriptional region, were enriched in the central region of TADs (**Fig. 5B and Supplementary Fig. S11C**). Then, we merged similar element types into nine broader categories. Furthermore, we defined 20 types of topological domains using the HMM. Based on the proportions of these nine categories within each domain type, we integrated the domains into six clusters (**Supplementary Fig. S12A**). Domains in clusters 1 and 6 showed high proportions of promoter/enhancer types and H3K36me3-dominant regions and had relatively fewer low-signal regions. Clusters 2 and 3 contained domains with high proportions of H3K27me3-dominant region and open chromatin, respectively. Domains in clusters 4 and 5 had a very low proportion of promoter/enhancer types and were predominantly characterized by low signal regions. (**Fig. 5C**). We found that cluster 5 had the highest proportion of domains, followed by clusters 1, 6, and 4, while clusters 2 and 3 were relatively less abundant (**Supplementary Fig. S12B**). The domains within these clusters show no significant differences in length, except for slightly larger domains in clusters 2 and 4 (**Supplementary Fig. S12C**). We also observed that domains in clusters 1 and 6 showed a higher proportion of compartment A and higher gene expression levels, while domains in clusters 4 and 5 showed the opposite trends (**Fig. 5D, E**). Moreover, domains in clusters 1 and 6 tended to replicate early and exhibited enhanced signals for H3K27ac and H3K36me3. Domains in cluster 2 showed intense signals for H3K27me3, while those in cluster 3 were characterized by marked DNase-seq signals. Domains in cluster 5 displayed stronger signals for H3K9me3 (**Supplementary Fig. S12D**). These results imply that domains in different clusters exhibit distinct chromatin states and biological properties. Additionally, using the Hidden Markov Model, we derived the transition probabilities between the 20 types of domains, which allowed us to group them into two sets with higher mutual transition probabilities within the same set (**Supplementary Fig. S12E**). The two distinct sets are primarily composed of domain clusters 1 and 5, corresponding to the most active and inactive domains, respectively (**Supplementary Fig. S12F**). It implies that the topological domains with similar characteristics tend to be clustered along the genome. Moreover, the active domain clusters exhibited stronger chromatin interactions, while the repressive domains predominantly interacted with other repressive ones (**Supplementary Fig. S12G**). It suggests that domains with similar properties also tend to aggregate in 3D space.

**Figure 5.**
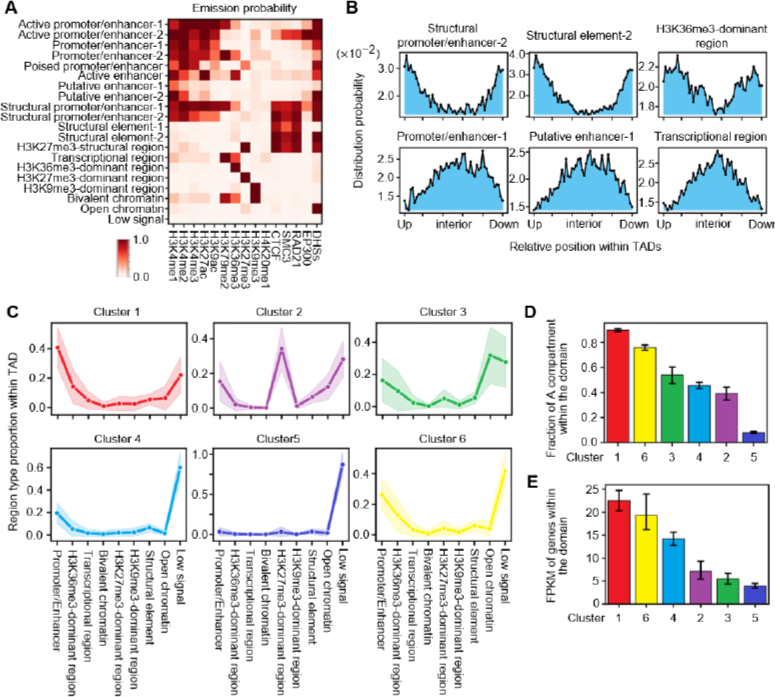
Functional annotations of TADs and the internal regions from a two-layer hidden Markov model. (**A**) Emission probability of 20 kinds of region-level states under multiple epigenomic modifications. (**B**) Distribution probabilities of relative positions for six region-level states within TADs and results of the remaining 14 states are shown in Supplementary Fig. S11C. (**C**) Six domain clusters with varying proportions across nine region types. (**D**) Fraction of compartment A within domains of six clusters. (**E**) Expression level of genes located in domains of six clusters.

### Deciphering TADs in the very noisy and sparse contact maps of 21 human tissues and cell lines

We applied TADGATE to a Hi-C dataset comprising 21 human tissues and cell lines (6) for data imputation and TAD identification. The Hi-C contact maps for many cell types exhibit high sparsity, making the topological structures too ambiguous to identify and hindering subsequent biological analysis (**Fig. 6A, B**). For cell types suffering from severe sparsity, such as Lung and Psoas, TADGATE can impute the sparse contact maps while enhancing the topological domains (**Fig. 6B**). For moderately sparse cell types like the hippocampus, TADGATE can denoise the contact maps, thereby making the topological domains clear. As for cell types with low sparsity and fine quality, e.g., the human embryonic stem cell line H1, TADGATE can maintain the topological structures while denoising the maps (**Supplementary Fig. S13A**). Compared to the clustering results based on the original contact maps, the TADGATE-imputed maps allow for better clustering of different cell types that are in line with the germ layer categories, such as ectoderm, mesoderm, endoderm, and embryos-derived cell lines. More specifically, similar tissues or cell types, such as Cortex and Hippocampus, and cardiac tissues like Aorta, Left Ventricle, and Right Ventricle, can also be effectively clustered together (**Fig. 6D and Supplementary Fig. S13B**).

**Figure 6.**
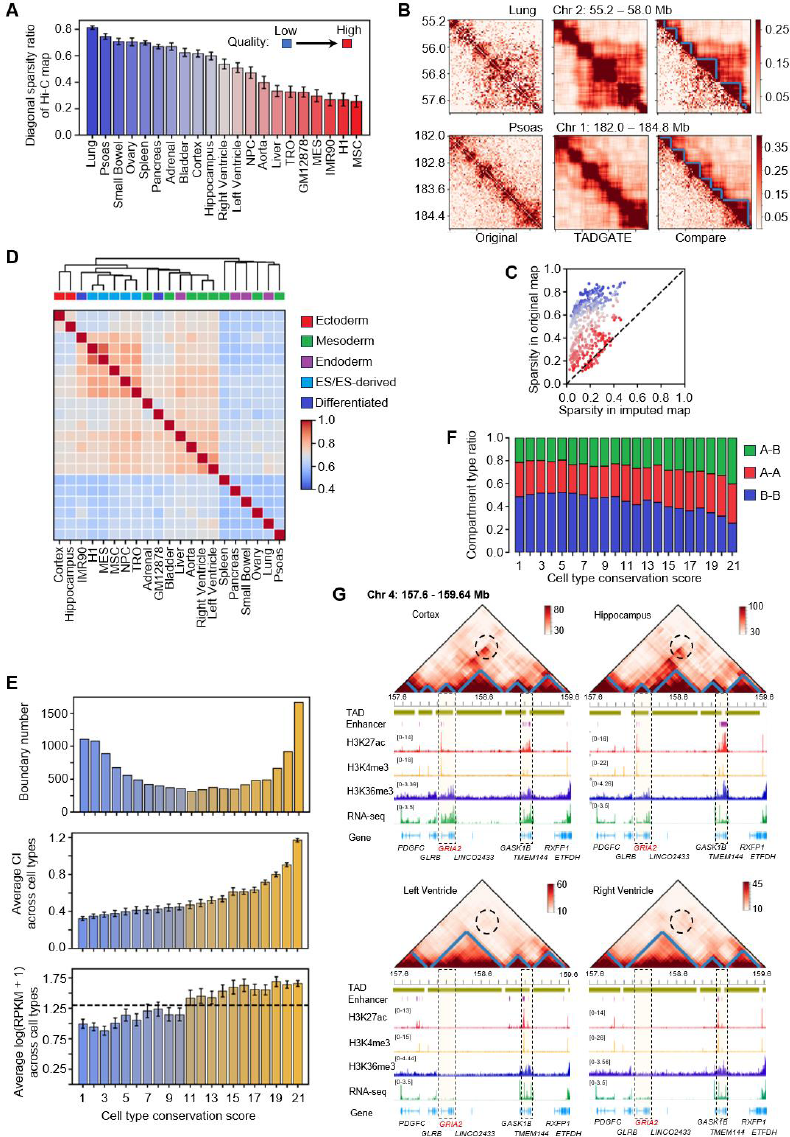
Analysis of Hi-C data of 21 human tissues and cell lines with TADGATE. (**A**) The diagonal sparsity ratio of Hi-C contact maps for all chromosomes in 21 human tissues and cell lines. Some cell types are abbreviated, such as Human Embryonic Stem Cell (H1), Mesendoderm (MES), Mesenchymal Stem Cell (MSC), Neural Progenitor Cell (NPC), and Trophoblast-like Cell (TRO). A high sparsity ratio represents poor quality of the contact map. (**B**) Two example regions to show the original Hi-C contact maps of lung and psoas, and the TADGATE-imputed contact maps and the corresponding TADs. In the third figure, the lower triangle represents the original map, while the upper triangle represents the TADGATE map. (**C**) Comparison of the diagonal sparsity ratios of the original maps and the TADGATE-imputed maps for all chromosomes in 21 tissues and cell lines. Each dot represents a chromosome and it is colored according to the cell type in (A). (**D**) Clustering results of all cell types based on the Spearman correlation coefficient of the Contrast Index with TADGATE-imputed contact maps for 21 tissues and cell lines. (**E**) The boundary number (top), the average contrast index across corresponding cell type (middle), and the average expression level of the nearby genes across all cell types (bottom) for core boundary regions with different cell type conservation scores. The dashed lines in the bottom figure represent the average gene expression at shuffled boundaries. (**F**) The compartment type ratios for boundary regions with different cell type conservation scores. (**G**) The comparison of TADGATE-imputed contact maps, chromatin topological domains, epigenetic signals, and RNA-seq signal around the gene *GRIA2* in the cortex, hippocampus, left ventricle, and right ventricle.

We defined some core boundary regions and assigned a cell type conservation score to each boundary, representing the count of cell types that exhibit the boundary (**Supplementary Fig. S13C, D**). TAD boundaries exhibited distinct conservation among different cell types, with over 50% of boundaries appearing in at least 11 cell types. But they also showed some cell or tissue specificity, with some boundaries appearing only in one or two cell types (**Fig. 6E**). The conserved boundaries among different cell types tended to have stronger insulation effects on chromatin interactions and the genes at conserved boundaries showed higher average expression levels across diverse cell types (**Fig. 6E**). Moreover, boundaries with varying conservation scores exhibited distinct compartmental environments. The probability of TAD boundaries being located between compartments of A-B or A-A type increased with a higher conservation score (**Fig. 6F**). The GO enrichment analysis showed that genes near conserved boundaries were often associated with fundamental biological processes, such as protein metabolic process, enzyme binding, and cytoplasmic composition (**Supplementary Fig. S13F**). However, we also found that genes at cell type-specific boundaries had lower average expression levels across all cell types (**Fig. 6E**). These genes didn’t show significantly higher expression ranks in the corresponding cell types (**Supplementary Fig. S14A**). However, the topological domains associated with these cell type-specific boundaries may contain genes related to the cell functions. For example, we observed that the genes located in domains around the Cortex or Hippocampus-specific boundaries were enriched in GO terms related to brain functions, such as Regulation of synapse organization and Axon guidance (**Supplementary Fig. S14B**). While genes in topological domains around boundaries specific to ES-derived cell lines were enriched in GO terms like Stem cell development and Embryonic organ development (**Supplementary Fig. S14C**). These results indicate that the organization of topological domains is highly related to the functions of tissues and cell types.

Moreover, we conducted pairwise comparisons between two tissues or cell types, revealing a significant negative correlation between the similarity of topological domains’ organization and the relative differences in gene expression (**Supplementary Fig. S14D**). With a comparison between the Hippocampus and Left Ventricle, as well as the Cortex and Right Ventricle, being examples, we found that the genomic regions with similar organization of topological domains showed smaller differences in gene expression. Conversely, the variation of topological domains may relate to the divergence in gene expression (**Supplementary Fig.S14E, F**). However, how topological domains affect gene expression may vary. For example, compared to the Left Ventricle and Right Ventricle, a unique topological domain was formed around the gene *GRIA2* in the Cortex and Hippocampus. The specific TAD interacted with another distant domain containing several enhancers or super-enhancers, which might enhance the expression of *GRIA2*. In contrast, in the Left or Right Ventricle, *GRIA2* resided in a large domain and didn’t show significant interactions with the distant domain although it also contained some enhancers (**Fig. 6G**). The expression level of *GRIA2* in the Cortex and Hippocampus was indeed higher than that in the ventricles (**Supplementary Fig. S15A**). Some studies have shown that *GRIA2* is closely related to brain function and correlated with many neurodevelopmental disorders (42, 43). By examining its expression levels in various human tissues (44), we observed that the gene *GRIA2* showed significant specific expression in the brain (**Supplementary Fig. S15D**). Therefore, our research provides a potential mechanism, from the perspective of topological domains, to understand the cell type-specific transcriptional regulation of the gene *GRIA2* in the brain, and it also suggests that chromatin can affect gene expression by forming unique TADs and shaping interactions between TADs.

Furthermore, some other examples also suggest that the difference in TAD boundaries may affect the gene expression in different cell types. For instance, the distributions of topological domains around the gene *PHYHIPL* in the Cortex, Hippocampus, and Left and Right Ventricles are very similar. *PHYHIPL* is located near a conserved TAD boundary in all cell types. However, the insulation effect of this TAD boundary in the Cortex and Hippocampus was weaker compared to the ventricles, which permitted the gene to interact more strongly with enhancers in the adjacent TAD. Consequently, this led to stronger expression of gene *PHYHIPL* in the Cortex and Hippocampus (**Supplementary Fig.S15B, C**). *PHYHIPL* also showed significant specific expression in brain tissues (**Supplementary Fig. S15E**). Although the exact function of the gene *PHYHIPL* in the human brain is not fully understood, studies have suggested its potential association with cerebellum-related disorders in mice (45). Therefore, further understanding of the transcriptional regulation of this gene is meaningful for subsequent research. Additionally, we found an event of TAD split near the genes *MYH6* and *MYH7* in the Left Ventricle and Right Ventricle compared to the Cortex and Hippocampus (**Supplementary Fig. S16B**). The genes *MYH6* and *MYH7* are located in a large topological domain in the Cortex and Hippocampus, whereas a new boundary forms near these genes in the ventricles and splits the large domain into two smaller ones. The distribution of enhancers in this region also changed. Correspondingly, the expression levels of *MYH6* and *MYH7* are higher in the ventricles (**Supplementary Fig. S16A, C**). These genes encode two kinds of myosin-heavy chains, which are major components of the thick filaments of the sarcomere in the mammalian heart (46). Both genes exhibit strong tissue-specific expression patterns in the heart (**Supplementary Fig. S16D, E**). The specific topological domains they reside in might explain their expression abundance. These two examples illustrate that changes in TAD boundary strength and the formation or absence of TAD boundary, which may cause TAD to split or merge, can also influence gene expression.

## Conclusions and Discussion

In this study, we achieve the smoothing, imputation, and topological domain enhancement of extremely sparse Hi-C contact maps with TADGATE for identifying accurate and robust TADs. The learned attention patterns reveal two types of units with distinct properties within TADs. We demonstrate the strong correlation between TADs and chromatin compartmentalization. The TAD boundaries in different compartmental environments showed varying insulation strength and diverse biological relevance. Detailed functional annotation of TADs and their internal regions reveals the distribution of structural and functional elements within TADs and different chromatin states of TADs, uncovering the aggregation phenomenon of TADs with similar states in both one-dimensional genome and three-dimensional space. TADGATE can be applied for cross-cell type TAD boundary comparison and reveal conserved and cell type-specific boundaries with distinct insulation strength and expression patterns. The imputed contact maps, with clearer topological domains, help to decipher cell type-specific gene regulation associated with chromatin topological structure alterations.

However, TADGATE provides imputed contact maps as a smoothed version of the original ones, which can facilitate the identification of TADs, especially in sparse and noisy contact maps. However, the smoothing effect may be inefficient in resolving finer chromatin structures, e.g., chromatin loops. In some cases, with high-quality contact maps, excessive smoothing could potentially obscure the depiction of chromatin loops. On the other hand, TADGATE aggregates the information of neighboring bins to reconstruct the short-range interactions, but it does not account for long-range chromatin interactions. Consequently, the TADGATE-imputed maps may not facilitate the study of large-scale chromatin structures, such as compartments or subcompartments.

There are potential directions for development in future research. With the continuous generation and accumulation of single-cell Hi-C data, which often suffer from increased sparsity, extending TADGATE to impute these sparser contact maps and identify TAD-like domains is a valuable direction. Additionally, recent advances in sequencing technologies, such as Pore-C (47) and HiPore-C (48), which can detect multi-way chromatin interactions, provide opportunities for further analysis of interactions between the structural and functional elements within TADs. It may facilitate our understanding of the more refined transcriptional regulatory processes within TADs. Besides, we assigned different categories and chromatin states to TADs. Future research on the interactions between TADs or discovering TAD communities with distinct properties is crucial for studying the overall organizational manners and functions of TADs.

## Materials and Methods

### Datasets

We used *in situ* Hi-C contact maps for the GM12878 and K562 cell lines (GEO accession: GSE63525) at 50 kb resolution and applied the ice-normalization to the map of each chromosome. We collected corresponding ChIP-seq, DNase-seq, Repli-seq, and RNA-seq data for these cell lines from the ENCODE project (49). We also got the chromatin states annotated by ChromHMM (36) and the subcompartments annotated by SNIPER (50). We also used the Hi-C contact maps and RNA-seq data for 21 human tissues and cell types (GEO accession: GSE87112). We constructed the contact maps at 40 kb resolution and performed the ice-normalization. We collected the corresponding ChIP-seq data for these tissues or cell types from the Roadmap project (51). All details of the data used in this study are reported in **Supplementary Table S1**.

### Methods TADGATE

#### Construction of the neighborhood graph

For the input iced-normalized chromatin contact map, where each row represents a genomic bin, TADGATE initially constructs a neighborhood graph based on a predefined radius. This graph reflects the 1D genomic proximity between bins, with each bin connected only to the upstream and downstream bins falling within the radius. The radius used in this study is 2.

#### Graph attention auto-encoder

Based on the neighborhood graph of genomic bins, we employed the graph attention layers in an auto-encoder to learn the latent embeddings of bins and reconstruct the chromatin contact map. The auto-encoder consists of an encoder and a decoder. The encoder takes the normalized interaction vector of each bin as input. Let *x_i_* be the normalized interaction vector of bin *i* and *L* represents the layer number of the encoder. By treating the interaction vectors as initial bin embeddings 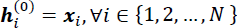, where *N* represents the total number of bins. The 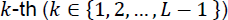 layer of the encoder is formulated as:

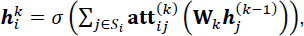

where 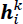 the intermediate embedding of bin *i* in layer *k*, ***W****_K_* is the trainable weight matrix, σ is the nonlinear activation function, *S_i_* is the neighbor set of bin *i* in the neighborhood graph (including bin *i* itself), and 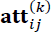 is the attention weight between bin *i* and bin *j* in the *k*-th graph attention layer. The last layer of the encoder adopts a fully connected manner: 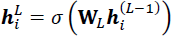. The low-dimensional embedding 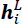 of each bin is the output of the encoder. On the flip side, the decoder *f_dec_* starts with the embedding 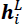 of each bin and reverses it into the reconstructed interaction vector, i.e., 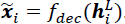. Our encoder employs one graph attention layer with 100 nodes and one fully connected layer with 50 nodes. The structure of the decoder is reversed. The objective of TADGATE is to minimize the reconstruction loss of the interaction vectors, 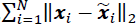. Since TADs are typically located near the diagonal of the contact map, we focus on the reconstruction errors specifically around the diagonal, with the default width being 10Mb.

The core part of TADGATE is the graph attention layer, which adaptively learns the topological similarity between neighboring bins. In the *k*-th encoder layer, the initial edge weight between bin *i* and its neighbor bin *j* is computed as follows:

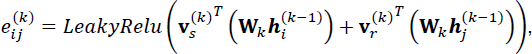

where 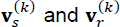 are the trainable weight vectors and *LeakyRelu* represents the activation function. Then, the edge weights are scaled as 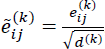, where *d*^(*k*)^ is the dimension of 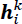. The scaled weights are normalized across all neighbors of bin *i* with a softmax function as follows:

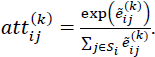

These learned attention weights are further used to update the latent embeddings of bins in the encoder and decoder.

#### Identification of TADs based on the output of graph attention auto-encoder

The output of the graph attention auto-encoder consists of three parts: the imputed Hi -C contact map, the latent embeddings of genomic bins, and the attention map. Combining the input contact maps, we integrate the four aspects to identify TADs (**Supplementary Fig. S1B**). First, for the original and imputed contact map, we calculate the contrast index (CI) and contrast *p*-value with a window size of 5 (**Supplementary Fig. S1A**). The CI peaks correspond to potential TAD boundaries, and the contrast *p*-values are used to assess whether the potential boundaries show a significant insulation effect on chromatin interactions. Second, we use Mclust (52) to cluster bins based on their latent embeddings. The optimal number of clusters is automatically selected using the BIC criterion from the Mclust package. We identify genomic bins with changing cluster labels compared to upstream or downstream bins as potential boundaries. Summing up the attention map by columns, we get the attention sum profile containing distinct peaks and valleys. Since we observed a correspondence between the attention valleys and TAD boundaries in the contact map, we also selected the attention valleys as potential boundaries. Collecting all potential boundaries from different sources, we employ a boundary voting strategy (31) to obtain boundary scores for each bin (**Supplementary Fig. S1B**). We then identify peaks in the boundary score profile and select the bins as true boundaries if they have contrast *p*-values below the given threshold for original or imputed contact maps. The threshold of contrast *p*-values used in this study is 0.05. Subsequently, we match adjacent boundaries and classify the intermediate region as domains, gaps, or boundary regions based on their average chromatin contacts.

### Comparison TADGATE with other TAD-calling methods

We compared TADGATE with four competing methods, i.e., DI, IS, TopDom, and GRNiCH. For all methods, we used the default or recommended parameter settings. However, when testing the performance of different methods on down-sampled data, we found that GRiNCH with default parameters failed to reconstruct clear TADs at most down-sampling ratios. Therefore, in the down-sampling analysis, we adjusted the expected length of TADs from 1Mb to 500kb in the default parameters of GRiNCH.

We compared the number and length of TADs identified by different methods in the Hi-C contact maps of GM12878. Besides, we treated each bin in the genome as a sample and used its interaction vector with other bins as the feature. Considering each TAD as a cluster of bins, we calculated the silhouette coefficient and Davies-Bouldin index to assess the TADs identified by each method. Both metrics are used to evaluate the effectiveness of clustering, and a higher silhouette coefficient or a smaller Davies-Bouldin index indicates better clustering results. We also compared the average enrichment levels of CTCF, RAD21, and SMC3 at TAD boundaries identified by different methods. We defined a unified boundary set based on the consensus among different methods (**Supplementary Fig. S1C**). In detail, we integrated the boundaries identified by five methods and assigned a boundary score to each bin through boundary voting to represent the number of methods that considered it as a boundary. We extracted contiguous bins with non-zero scores to form boundary regions. Subsequently, we examined whether each boundary region exhibited insulation effects on chromatin contacts and enrichment effects of CTCF. We used the filtered ones as the high-confident TAD boundaries. By examining whether the boundaries identified by each method overlap with these high-confident ones, we can determine the number of true boundaries (TB), false boundaries (FB), and missed boundaries (MB), and then calculate the corresponding precision, recall, and F1-score.

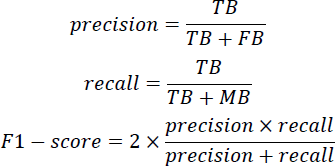

On the other hand, both TADGATE and GRiNCH can impute and denoise the chromatin contact maps. Therefore, we designed the down-sampling experiment based on the contact maps of chromosomes 2 to 5 of the K562 cell line. We randomly sampled a certain proportion of sequencing reads from the complete dataset and constructed the corresponding down-sampled contact maps. Subsequently, we applied five TAD-calling methods to these down-sampled maps. We used the HiCrep score, peak-to-signal ratio (PSNR), and structure similarity (SSIM) to evaluate the similarity between the contact maps with 100% reads and the imputed maps produced by TADGATE or GRiNCH. We also obtained the unified boundary set in the contact maps with 100% reads and calculated the precision, recall, and F1-score for all five methods at different down-sampling ratios. We used the Jaccard index to assess the overlap of boundaries identified by each method in the full-reads and down-sampled maps,

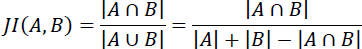

where 0 ≤ *JI(A, B)* ≤ 1. A and B represent two boundary sets. If the distance between two boundaries doesn’t exceed 1 bin, they will be determined as the shared ones between boundary sets.

### Analysis of TADs and chromatin compartment

We identified the A/B compartment at 100kb resolution using the PCA-based method (4) and downloaded the subcompartment annotations of GM12878 and K562 obtained by SNIPER (50). When defining the compartment purity of TADs, if a compartment or subcompartment comprises more than 10% of the TADs, we consider that the TAD contains this compartment or subcompartment. We also examined the distance between compartment or subcompartment switch points and TAD boundaries, employing shuffled TADs as a control. For each TAD boundary, we categorized them into three types: A-A, A-B, and B-B, based on the compartment types on either side of them. We compared the epigenomic modifications around the boundaries of different compartment types and subtracted the mean value from the signal vector of each boundary to eliminate the influence of genomic position bias. For enrichment analysis of repeat elements in the TAD boundary, we shuffled all TADs 100 times and calculated the number of repeat elements in identified and random boundaries, respectively. We then calculated the enrichment z-score of repeat elements in the boundary based on the mean and variance of the elements in random regions. Hence, a z-score above one indicates the repeat element is enriched in the boundary region.

### Comparison of TAD boundaries between GM12878 and K562 in terms of compartmental type

We identified TAD boundaries separately in GM12878 and K562 using TADGATE. Then, we categorized GM12878 or K562 boundaries into three types: A-A, A-B, and B-B, respectively. When comparing the boundary between GM12878 and K562, those with a distance of one bin or less were considered position-conserved. Among these position-conserved boundaries, we further classified them into compartment-type stable and transitioned boundaries based on their compartment types in both cell types. We applied the logarithmic transformation to the contrast index and calculated the z-score for each chromosome in both cell lines (**Supplementary Fig. S10A**). Thus, the normalized contrast index was comparable between the two cell types. Finally, by employing Fisher’s exact test, we examined whether more boundaries were associated with changes in the normalized contrast index among the compartment-type transitioned boundaries relative to the compartment-type stable boundaries.

### Functional annotation of TADs and the internal regions

We employed a two-layer Hidden Markov Model (41) combined with a set of epigenetic markers to annotate the TADs identified by TADGATE in GM12878 and their internal regions. We first divided each TAD into 50 regions of equal size. We checked whether each region contained the signal peaks for each epigenetic marker. If so, the region was assigned a score of 1, otherwise 0. After examining all markers, we obtained binary observation features for each region. Next, we trained a two-layer Hidden Markov Model with region and TAD as two hierarchical layers. The model required pre-specified hidden state numbers for regions and TADs. We tried different numbers of states and selected 20 as the optimal number according to the average explained variance for all markers (**Supplementary Fig. S11A**). The explained variance is defined as the rate of changes in the variance of all bins before and after obtaining the element-type label or the domain-type label for each epigenetic marker.

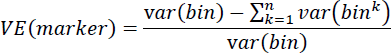

Where *var(bin)* represents the variance of all bins along the chromosome and *var(bin^k^)* represents the variance of bins belonging to the *k*-th type of region or domain. Thus, we classified TADs and the regions within TADs into 20 types. For the region-level types, we manually annotated them based on the emission probability matrix output by the model, such as Active promoter/enhancer, Structural element, and H3K27me3-dominant region (**Fig. 5A and Supplementary Fig. S11B**). This model also outputs a transition probability matrix between the 20 domain types (**Supplementary Fig. S12E**). To better annotate the domain types, we merged similar types among the 20 region types to obtain nine broader categories. Subsequently, we examined the proportions of the nine region categories among the 20 domain types and used this information to merge the 20 domain types into six clusters (**Fig. 5C and Supplementary Fig. S12A**). We analyzed the signals of Repli-seq, DNase-seq, and the ChIP-seq for H3K27ac, H3K36me3, H3K27me3, and H3K9me3 within the domains of each cluster. We divided each domain into 100 intervals and calculated the average value of each kind of signal for each interval (**Supplementary Fig. S12D**). We also compared the chromatin interactions between domains in each cluster. We first normalized the Hi-C contact maps and then calculated the z-score at each distance to eliminate the influence of genomic distance on the strength of chromatin interactions. Then, we examined the average chromatin interaction z-score between all domains from a pair of clusters (**Supplementary Fig. S12G**).

### Clustering of 21 human tissues and cell lines

The contrast index depicts the distribution of TADs in chromatin contact maps. We calculated the contrast index for all chromosomes in each cell type based on the original and the TADGATE-imputed maps. Then, for each cell type, we concatenated the vectors of all chromosomes and clustered the cell types based on the combined vector (**Fig. 6D and Supplementary Fig. S13B**).

### Defining the core boundary regions with conservation scores across 21 cell types

First, we used TADGATE to identify TADs in chromatin contact maps of 21 cell types. Then, we integrated the boundaries from all cell types and assigned a boundary score to each bin along the genome, representing the times it served as a TAD boundary in one cell type. Next, we extracted consecutive bins with non-zero scores to form boundary regions. For each boundary region, if there were multiple peaks with scores greater than five, we divided the region into several subregions, using the midpoints between these peaks as the split points. We selected the bin with the highest score as the core boundary region within each subregion. If the adjacent bins to the core had non-zero scores, we added them to the core boundary region. The length of each core boundary region did not exceed three bins. Then, we examined the contrast p-values of each core boundary region in the original and the TADGATE-imputed maps for each cell type. If there were bins with *p*-values below the threshold, we considered this core boundary region serving as a TAD boundary in that cell type, and the conservation score increased by one (**Supplementary Fig. S13C, D**). Therefore, the range of boundary conservation scores is from 1 to 21, and the boundary regions with high scores represent conserved boundaries across different cell types (**Fig. 6E, F**).

### Gene set expression rank analysis for cell type-specific boundaries

We defined boundary regions with conservation scores of 1 as cell type-specific boundaries and extracted the genes within them. We conducted expression rank analysis (6) on these gene sets to determine if they exhibit higher expression in specific cell types. First, we obtained the log(RPKM+1) for all genes. Then, we removed genes with zero expression across all cell types. For each gene, we normalized its expression by dividing the sum across all cell types and obtained the relative expression level in each cell type. Within each cell type, based on the relative expression of all genes, we assigned the expression rank to each gene, where higher relative expression values corresponded to higher ranks. For the gene set corresponding to cell type-specific boundaries, we can compute their average expression rank in any cell type. We randomly select an equal number of boundaries, extract their genes, and calculate the average rank in the same cell type. When the rank of the specific gene set is significantly higher than that of the random gene set, it indicates that these genes have specific higher expression levels in the current cell type. However, we found that the genes associated with cell type-specific TAD boundaries did not exhibit higher expression in the corresponding cell type (**Supplementary Fig. S14A**).

### Pairwise comparison between two cell types for topological domains and gene expression

We divided the genome into 2Mb intervals and calculated the Spearman Correlation Coefficient (SCC) for each interval between the contrast index profiles of the two cell types. A higher correlation coefficient indicates a higher similarity in the distribution of topological domains between the two cell types. Additionally, we calculated the average relative expression difference for all genes within each interval, defined as the difference in expression levels dividing the larger one of the two expression values, ranging from 0 to 1. A higher value indicates more distinct differences in gene expression between the two cell types. Furthermore, we could calculate the Spearman correlation coefficient between the similarity of the topological structures and the relative expression differences for all intervals. A negative correlation coefficient suggests that as the similarity of the topological structures increases, the differences in gene expression between the two cell types decrease (**Supplementary Fig. S14D-F**).

### Software availability

The TADGATE source code is available on GitHub (https://github.com/zhanglabtools/TADGATE).

## Acknowledgments

This work has been supported by the National Key Research and Development Program of China [No. 2019YFA0709501 to S.Z.], the National Natural Science Foundation of China [Nos. 62173271, 61873202 to S.W.Z.; 32341013, 12326614, 12126605 to S.Z.], the R&D project of Pazhou Lab (Huangpu) [No. 2023K0602 to S.Z.], and the CAS Project for Young Scientists in Basic Research [No. YSBR-034 to S.Z.].

