## Supplemental Figures for "Uncovering topologically associating domains from three-dimensional genome maps with TADGATE"

### Supplementary Figures

**Fig. S1.** The process of TAD identification by TADGATE and the process to get a unified boundary set.

**Fig. S2**. Comparison of different TAD-calling methods with Hi-C data of GM12878 cell line.

**Fig. S3**. Comparison of different TAD-calling methods with down-sampled Hi-C data of K562 cell line.

**Fig. S4.** TADs identified by TADGATE, GRiNCH, TopDom, IS and DI in contact maps at different down-sampling ratios.

**Fig. S5**. Evaluation of the boundary enhancement effect for TADGATE and GRiNCH.

**Fig. S6**. Relationship of attention profiles and domain structures.

**Fig. S7**. Analysis of epigenomic markers and chromatin states of attention peaks and valleys.

**Fig. S8**. The relationships between TADs and chromatin compartmentalization.

**Fig. S9**. Analysis of TAD boundaries and epigenomic modifications.

**Fig. S10**. The comparison of compartmental types of TAD boundaries between GM12878 and K562.

**Fig. S11**. The types of structural and functional elements within TADs and their distribution in TADs.

**Fig.S12.** Define the types of domain clusters and analyze their biological characteristics.

**Fig. S13**. Analysis of Hi-C data of 21 human tissues and cell lines with TADGATE.

**Fig. S14**. Analysis of genes around cell-type specific boundaries and the correlation between chromatin topological similarity and difference of gene expression.

**Fig. S15**. Analysis of the expression of genes *GRIA2* and *PHYHIPL* and the nearby structures of topological domains in different cell types.

**Fig. S16**. Analysis of the expression of genes *MYH6* and *MYH7* and the nearby structure of topological domains in different cell types.

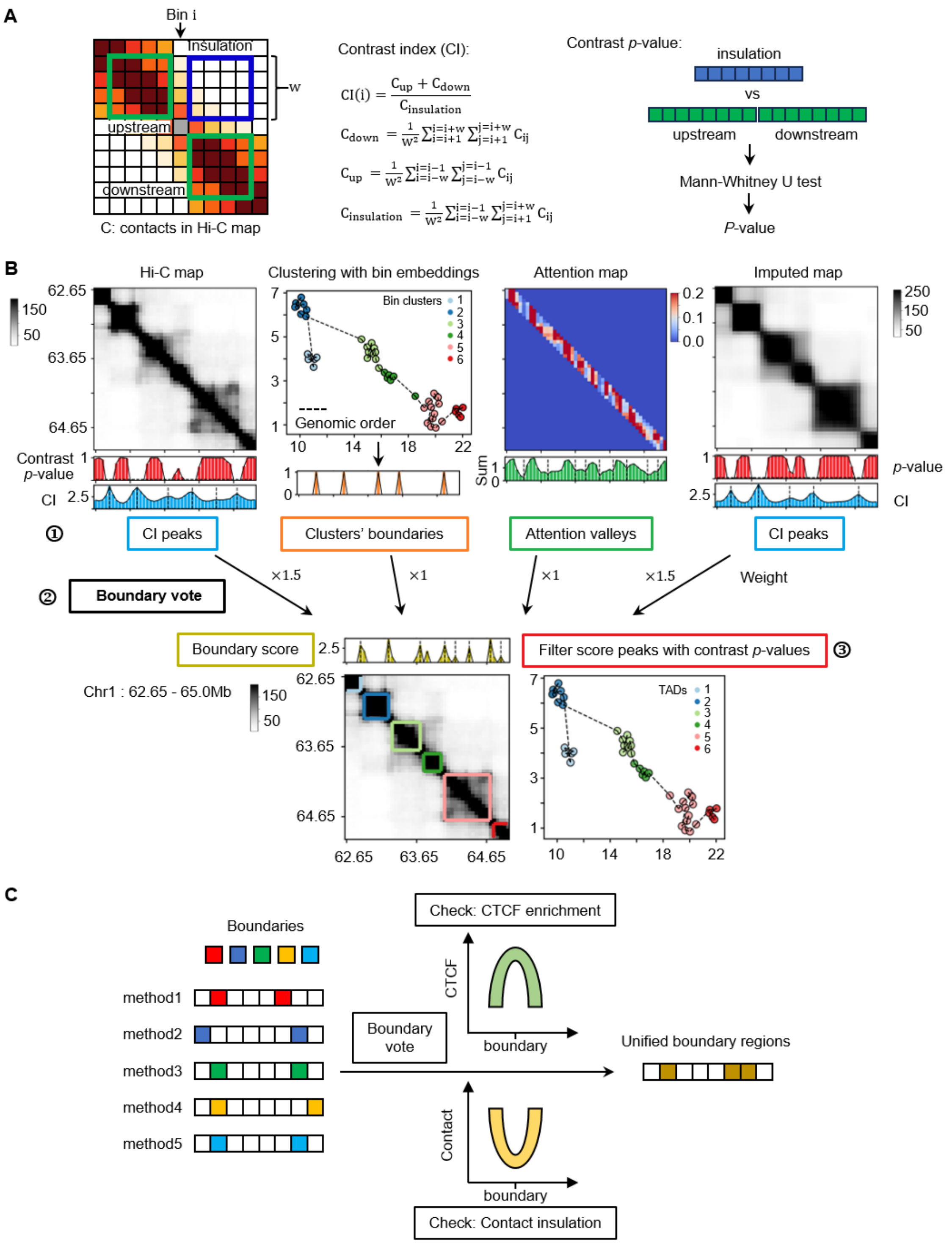


$$CI(i) = \frac{C_{up} + C_{down}}{C_{insulation}}$$

$$C_{down} = \frac{1}{w^2}\sum_{i=i+1}^{i=i+w}\sum_{j=i+1}^{j=i+w} C_{ij}$$

$$C_{up} = \frac{1}{w^2}\sum_{i=i-w}^{i=i-1}\sum_{j=i-w}^{j=i-1} C_{ij}$$

$$C_{insulation} = \frac{1}{w^2}\sum_{i=i-w}^{i=i-1}\sum_{j=i+1}^{j=i+w} C_{ij}$$

**Fig. S1. The process of TAD identification by TADGATE and the process to get a unified boundary set.** (**A**) Illustration of how to calculate contrast index (CI) and contrast *p*-value. (**B**) Illustration of the method to identify TADs based on the output of TADGATE. TADGATE obtains the boundary scores for each bin based on four signal sources. The contrast indexes and *p*-values of the original Hi-C map and the TADGATE-imputed map are first calculated and the corresponding CI peaks are identified. Bins are clustered based on the embeddings learned by TADGATE and the bins with changing cluster labels when compared to the upstream or downstream bins are identified. The attention valleys in the attention sum profile are also identified. These bins represent the candidate positions of TAD boundaries, which are then subjected to weighted voting to obtain boundary scores. The peaks in boundary score distribution are identified and then filtered with contrast *p*-values to get the final boundaries. (**C**) The process of constructing the unified boundary set of boundaries identified by different methods. The unified boundaries refer to bins with non-zero scores after boundary voting, and they are required to demonstrate both CTCF enrichment and insulation effects on chromatin contacts.

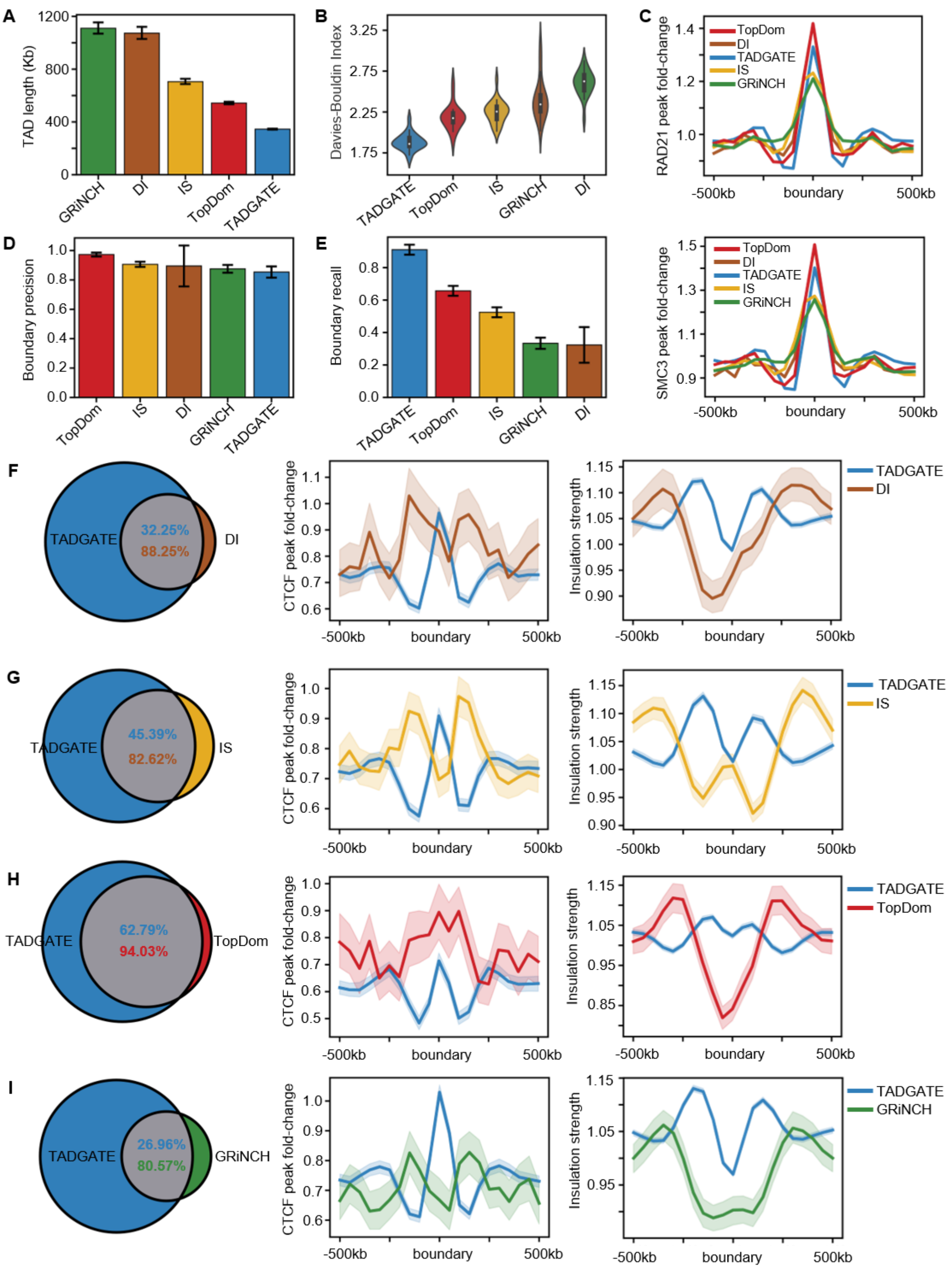


**Fig. S2**. **Comparison of different TAD-calling methods with Hi-C data of GM12878 cell line.** (**A**) Lengths of TADs identified by TADGATE and other four methods in chromosomes 1-22 and X of the GM12878 cell line. (**B**) Davies-Bouldin index of TADs identified by various methods on each chromosome, with each bin treated as a sample and a TAD as the cluster of corresponding bins. (**C**) RAD21 and SMC3 binding profiles around the TAD boundaries identified by different methods. (**D**) Precision of boundaries identified by different methods on each chromosome. (**E**) Recall of boundaries identified by different methods on each chromosome. (**F-I**) Pair-wise comparison of boundaries identified by TADGATE and other methods. The overlaps of boundaries are shown. The CTCF binding and insulation strength profiles around the unique boundaries of each method are shown.

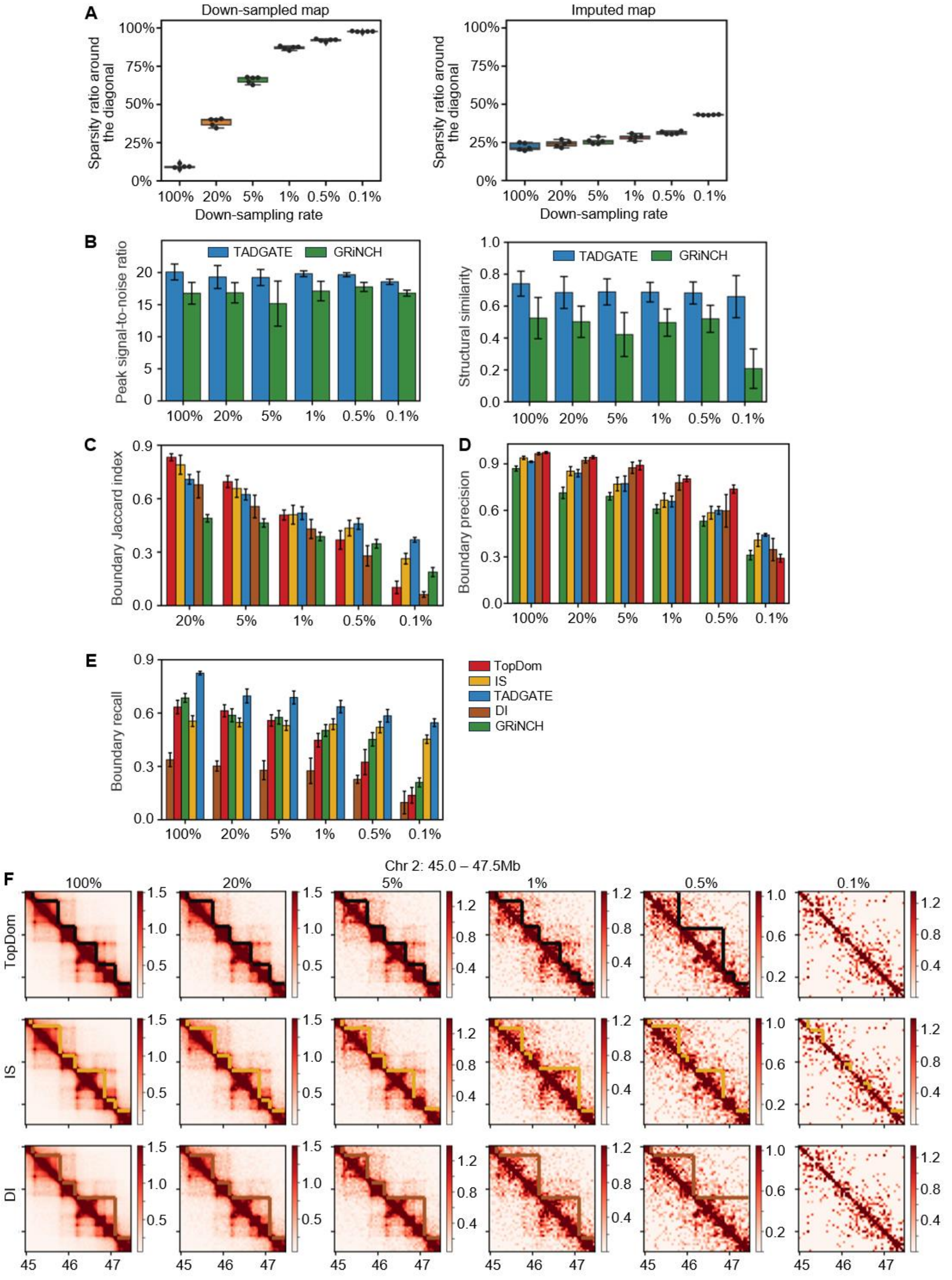


**Fig. S3**. **Comparison of different TAD-calling methods with down-sampled Hi-C data of K562 cell line.** (**A**) Sparsity ratio of chromatin contacts of the 10-Mb range around the diagonal of the down-sampled Hi-C contact maps and the maps imputed by TADGATE. The contact maps imputed by GRiNCH contain no empty values, i.e., 0% sparsity ratio, and its results are not shown. (**B**) Peak signal-to-noise ratio and structural similarity between the Hi-C contact map with 100% reads and the maps imputed by TADGATE or GRiNCH at different down-sampling ratios. (**C**) Jaccard index of boundaries identified in the Hi-C contact map with 100% reads and the down-sampled maps for each method. (**D and E**) Boundary precision and recall of boundaries identified by each method in contact maps at different down-sampling ratios. (**F**) TADs identified by TopDom, IS, and DI in contact maps at different down-sampling ratios, see Figure 2G.

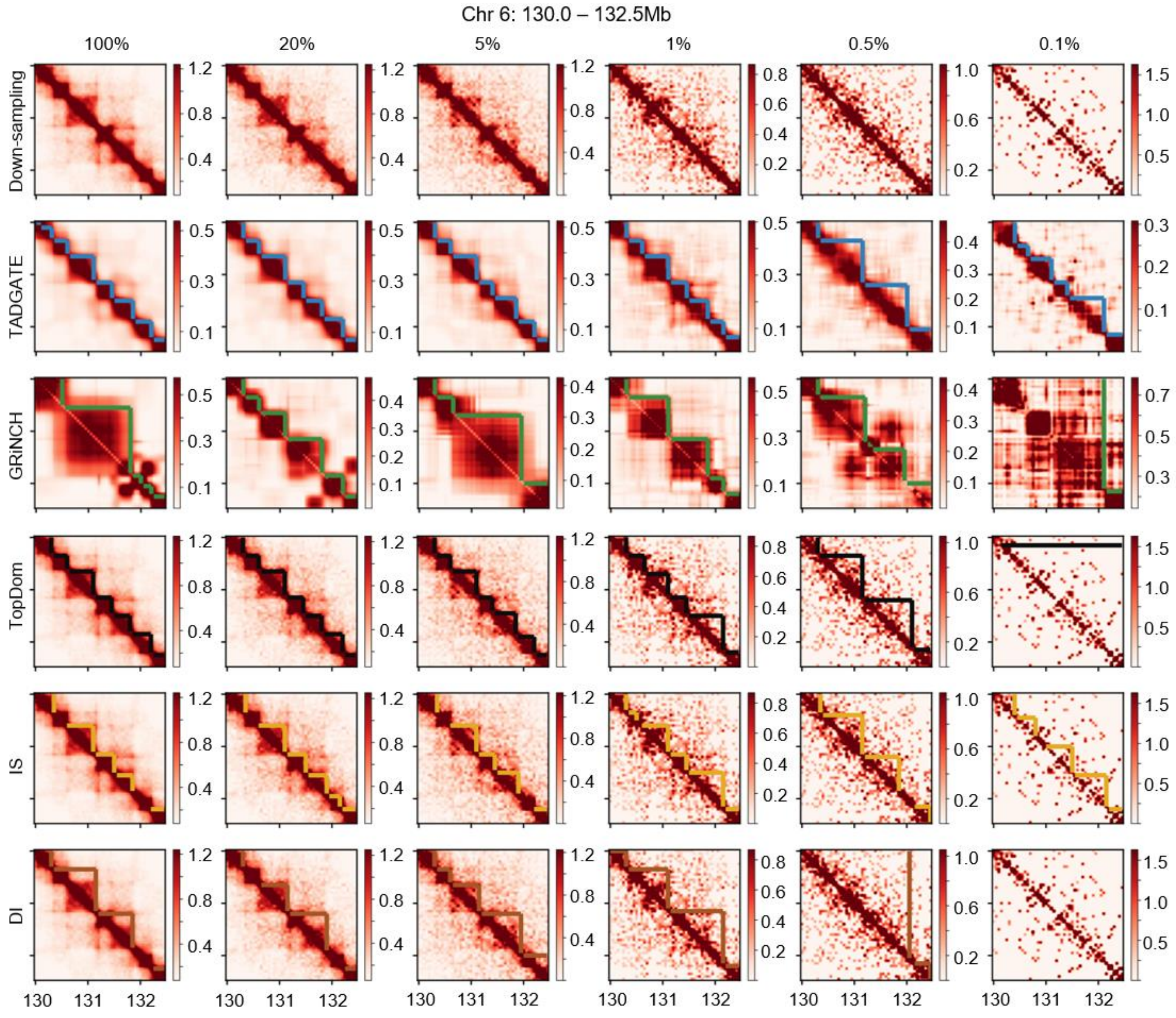


**Fig. S4.** TADs identified by TADGATE, GRiNCH, TopDom, IS, and DI in contact maps at different down-sampling ratios.

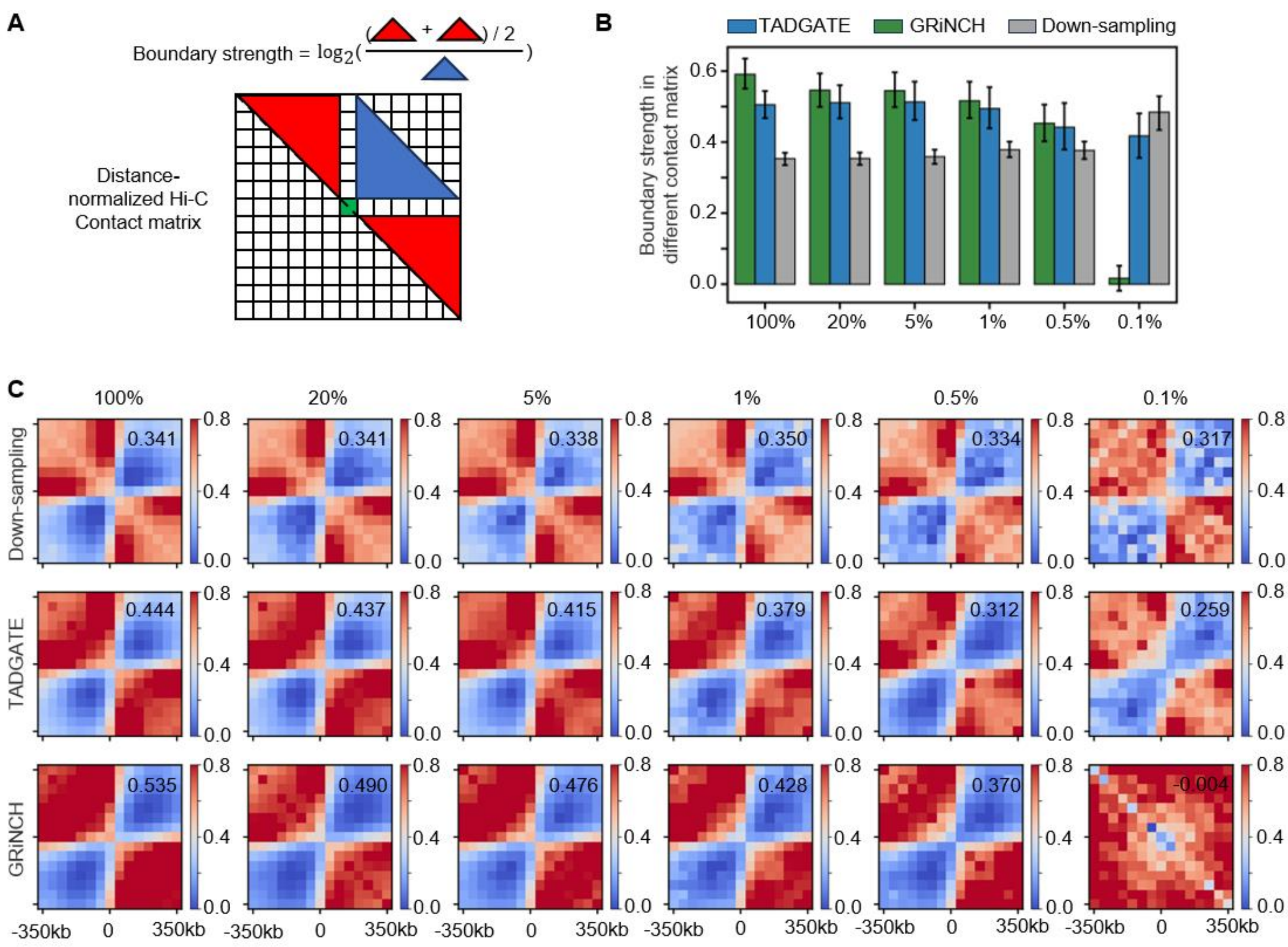


**Fig. S5**. **Evaluation of the boundary enhancement effect for TADGATE and GRiNCH.** (**A**) Illustration of the calculation of boundary strength. (**B**) The strength of boundaries from the unified sets within the down-sampled Hi-C contact maps and maps imputed by TADGATE or GRiNCH. (**C**) Aggregated maps around the boundaries from the unified set in down-sampled Hi-C contact maps and maps imputed by TADGATE or GRiNCH. The boundary strength in the aggregated map is displayed in the top-right corner.

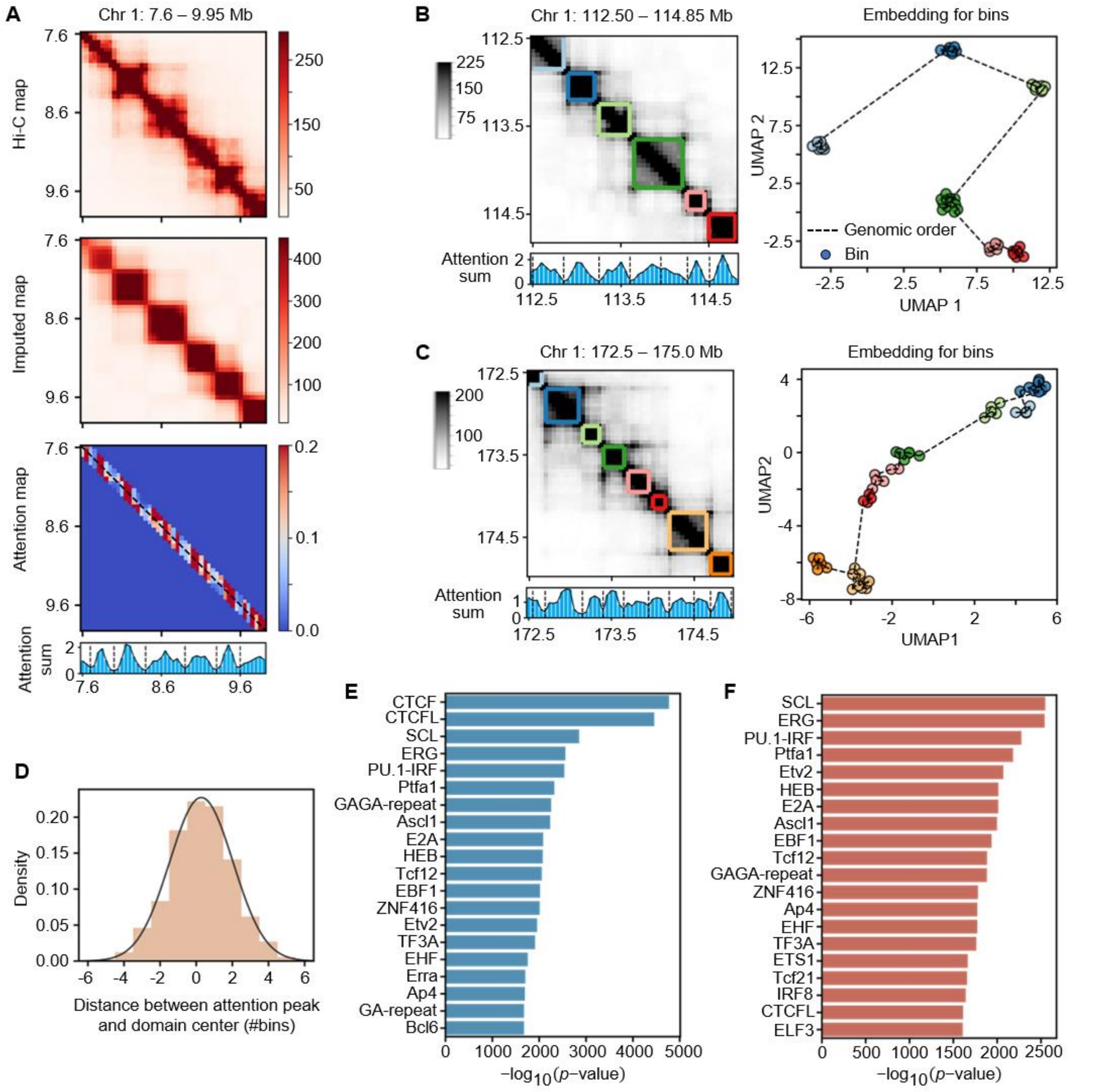


**Fig. S6**. **Relationship of attention profiles and domain structures.** (**A**) A case region to show the input Hi-C contact map, the TADGATE-imputed map, the neighborhood attention map, and the attention sum profile. (**B and C**) Hi-C contact maps of two representative regions with the TADs identified by TADGATE. The corresponding profile of attention sum is shown below, and dashed lines mark the attention valleys. The UMAP plot on the right is generated based on the embedding vectors of bins obtained from TADGATE, with bins colored according to the TADs they belong to. (**D**) Position distribution of attention peaks within domains. (**E and F**) TF motifs enriched in attention valleys (E) and attention peaks (F).

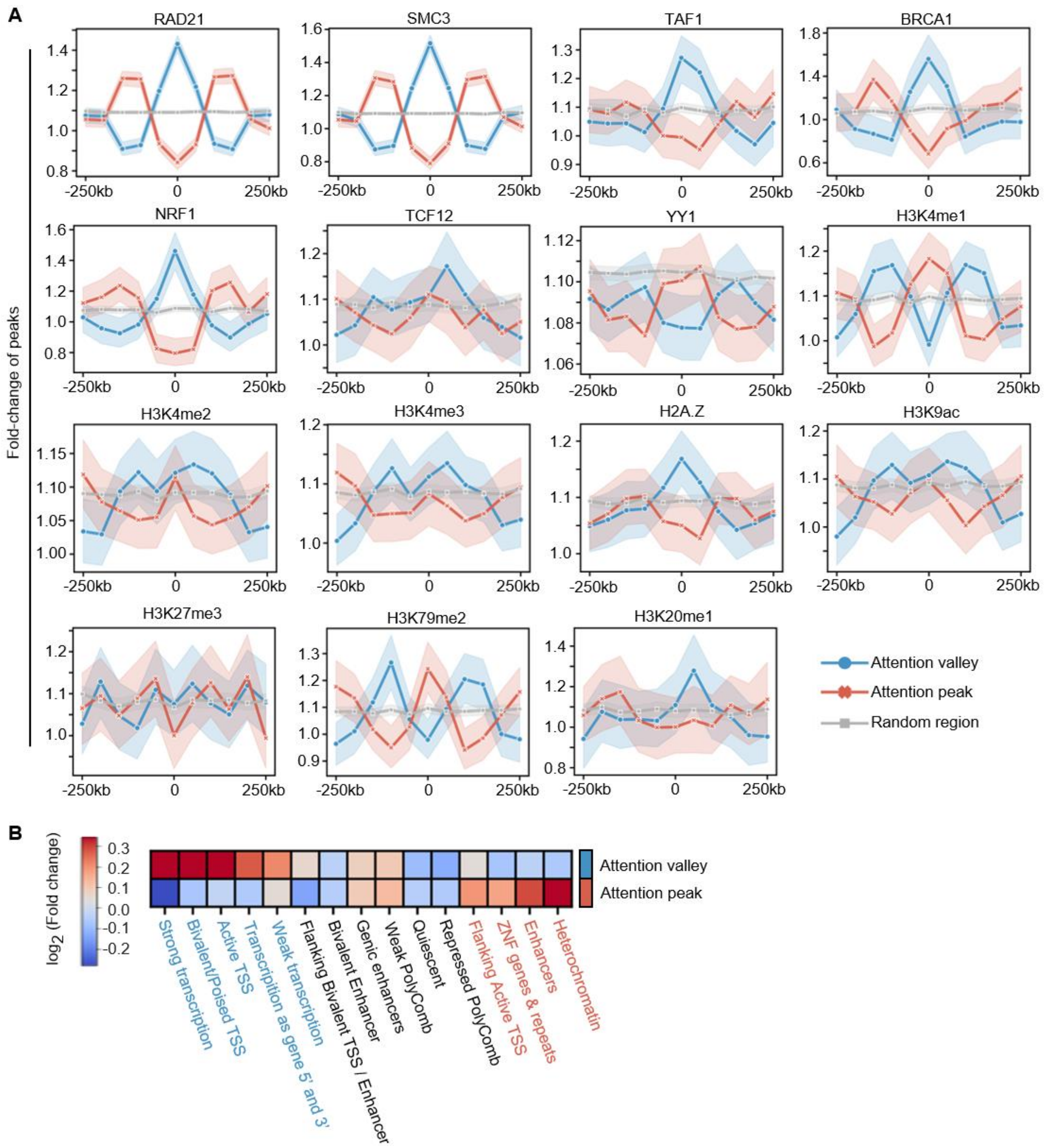


**Fig. S7**. **Analysis of epigenomic markers and chromatin states of attention peaks and valleys.** (**A**) Profiles of some transcription factors, histones, and DHSs around attention peaks or valleys. The shaded area represents the 95% confidence interval in bootstrap. See Figure 3G. (**B**) Enrichment of 15 chromatin states in attention valleys and peaks. The top enriched states are marked by blue for attention valleys and red for attention peaks.

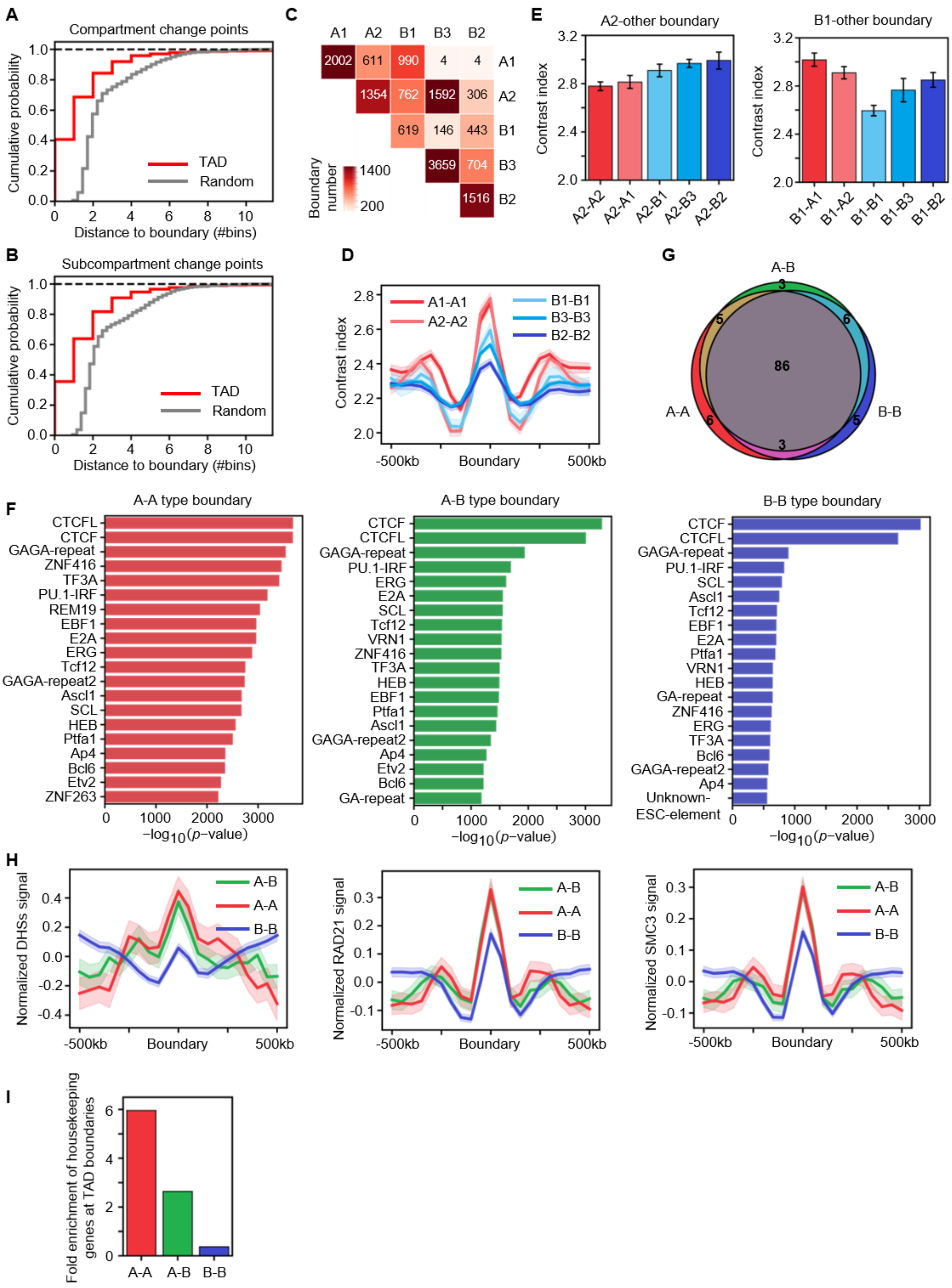


**Fig. S8**. **The relationships between TADs and chromatin compartmentalization.** (**A and B**) The accumulative probability distribution of the distance between the A/B compartment or subcompartment switch points and the nearest TAD boundaries or randomly shuffled boundaries. The Kolmogorov-Smirnov test is performed and both conditions show a *p*-value < 0.0001. (**C**) The number of TAD boundaries between different combinations of subcompartments. (**D**) The comparison of the contrast index of TAD boundaries between the same subcompartment types. (**E**) The contrast index of TAD boundaries between A2 type or B1 type and other subcompartments. (**F**) The top 20 enriched TF motifs in the three types of boundaries. (**G**) The overlap among the top 100 enriched TF motifs in the three types of boundaries. (**H**) The normalized signals of DHSs, RAD21, and SMC3 around the three types of boundaries. (**I**) The fold enrichment of housekeeping genes located at the three types of boundaries.

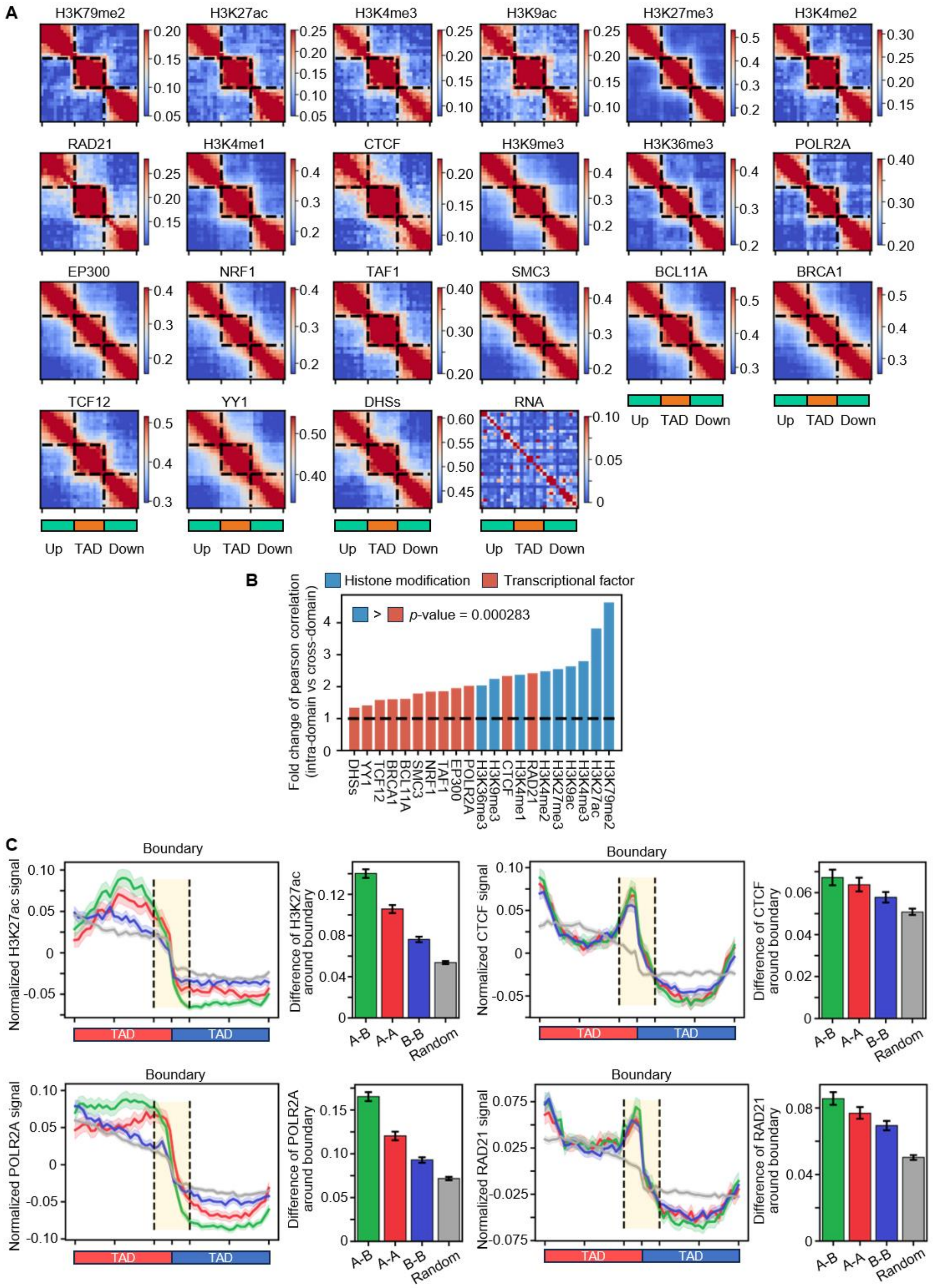


**Fig. S9**. **Analysis of TAD boundaries and epigenomic modifications. (A)** Pearson correlation coefficients (PCC) for the distribution of various epigenomic modifications between regions within TADs and upstream or downstream. (**B**) Fold-change of average PCC within TAD and average PCC between TAD and upstream or downstream regions. Single-end Mann-Whitney U tests are performed. (**C**) The profile of H3K27ac, POLR2A, CTCF, and RAD21 around TAD boundaries between different combinations of compartments. For each boundary, the direction of the profile is adjusted to keep a higher signal in the left TAD than the right one. The bar plot on the right side also exhibits the difference between the average signals within the two TADs.

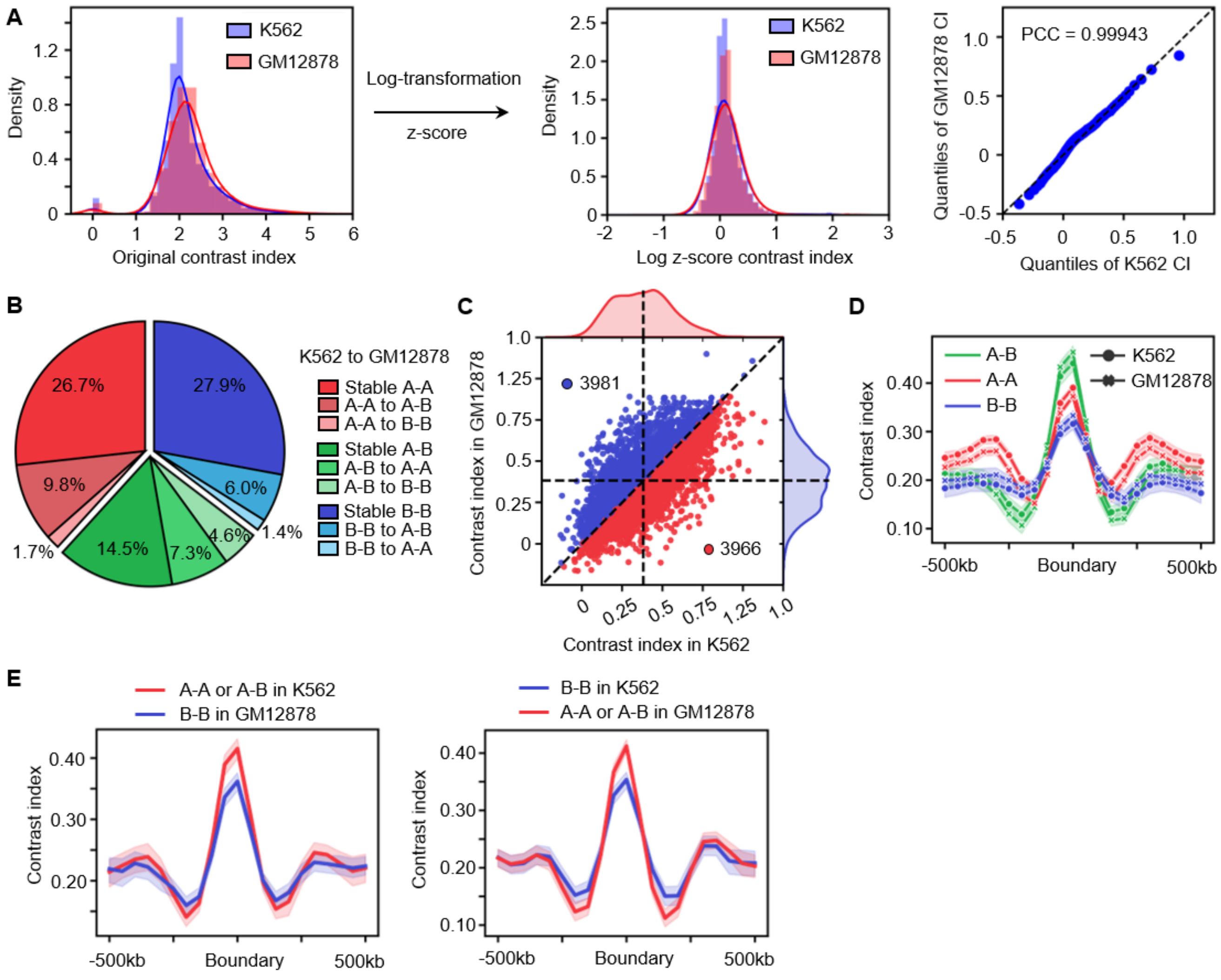


**Fig. S10**. **The comparison of compartmental types of TAD boundaries between GM12878 and K562.** (**A**) The probability density distributions of the original contrast index and the contrast index after logarithmic z-score transformation for Chromosome 2 in GM12878 and K562. The plot on the right shows the quantiles of the contrast index after transformation in GM12878 and K562, respectively. The Pearson correlation coefficient between these quantiles is also shown. (**B**) The proportion of different compartment-type transitions of the position-conserved TAD boundaries between K562 and GM12878. (**C**) The distribution of contrast index of TAD boundaries that exhibit conserved position and the same compartment types between GM12878 and K562. The boundaries with higher contrast index in each cell line are marked with red or blue, and the corresponding boundary numbers are also shown. The vertical and horizontal dashed lines indicate the mean contrast index of these boundaries in GM12878 and K562. (**D**) The profiles of contrast index around certain TAD boundaries in GM12878 and K562. These boundaries have conserved positions and the same compartment types between GM12878 and K562. (**E**) The profiles of contrast index around certain TAD boundaries in GM12878 and K562. These boundaries have conserved positions but different compartment types between GM12878 and K562.

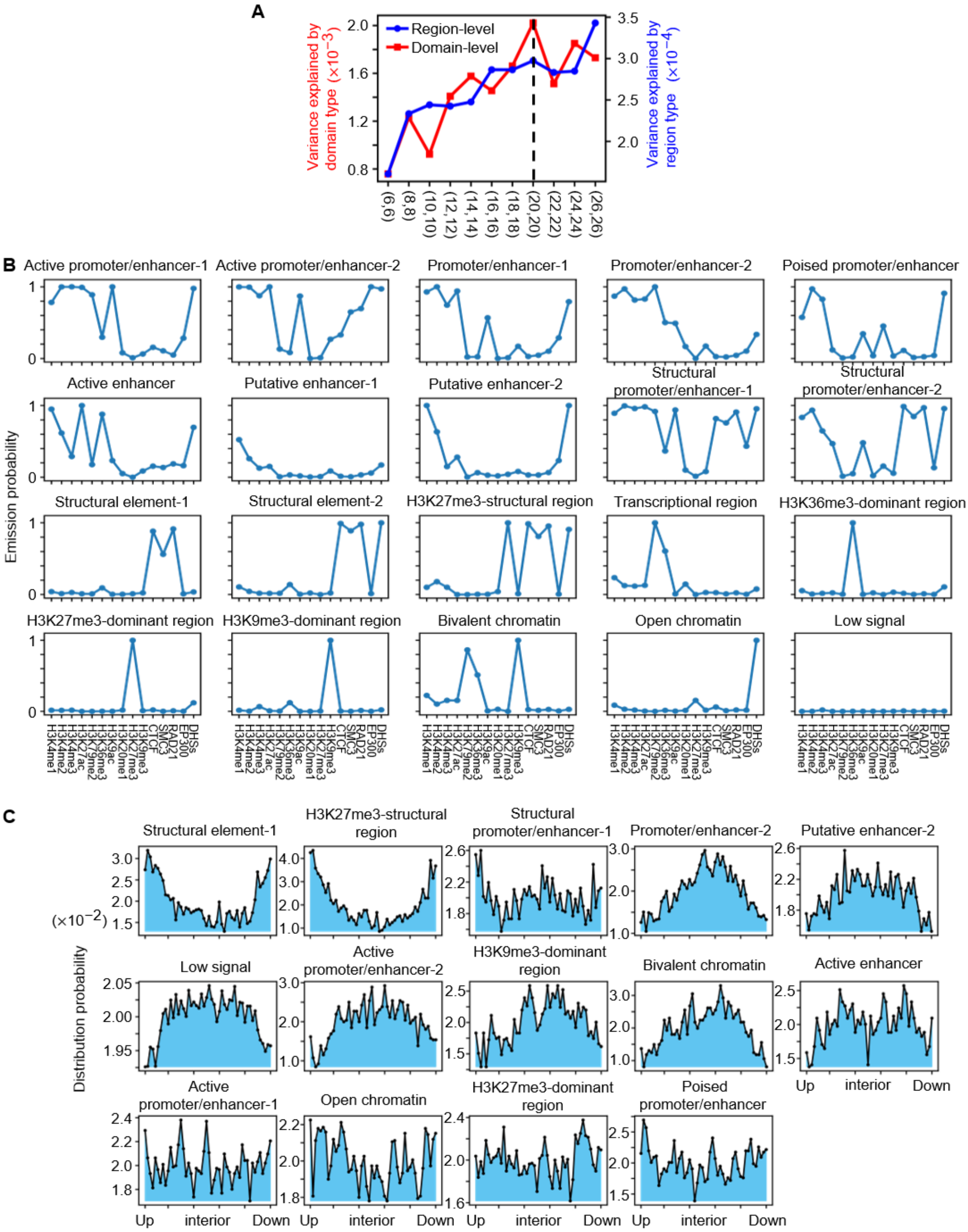


**Fig. S11**. **The types of structural and functional elements within TADs and their distribution in TADs.** **(A)** The average explained variance under different numbers of hidden states for domain-level and region-level in the two-layer Hidden Markov Model. The variance explained at the domain level is shown on the red axis, while the variance explained at the region level is shown on the blue axis. (**B**) The emission vector of each structural or functional element across multiple epigenomic markers, corresponding to each row of the emission matrix in Fig 5A. (**C**) Distribution probabilities of relative positions for 14 element-level states within TADs. See Fig. 5B.

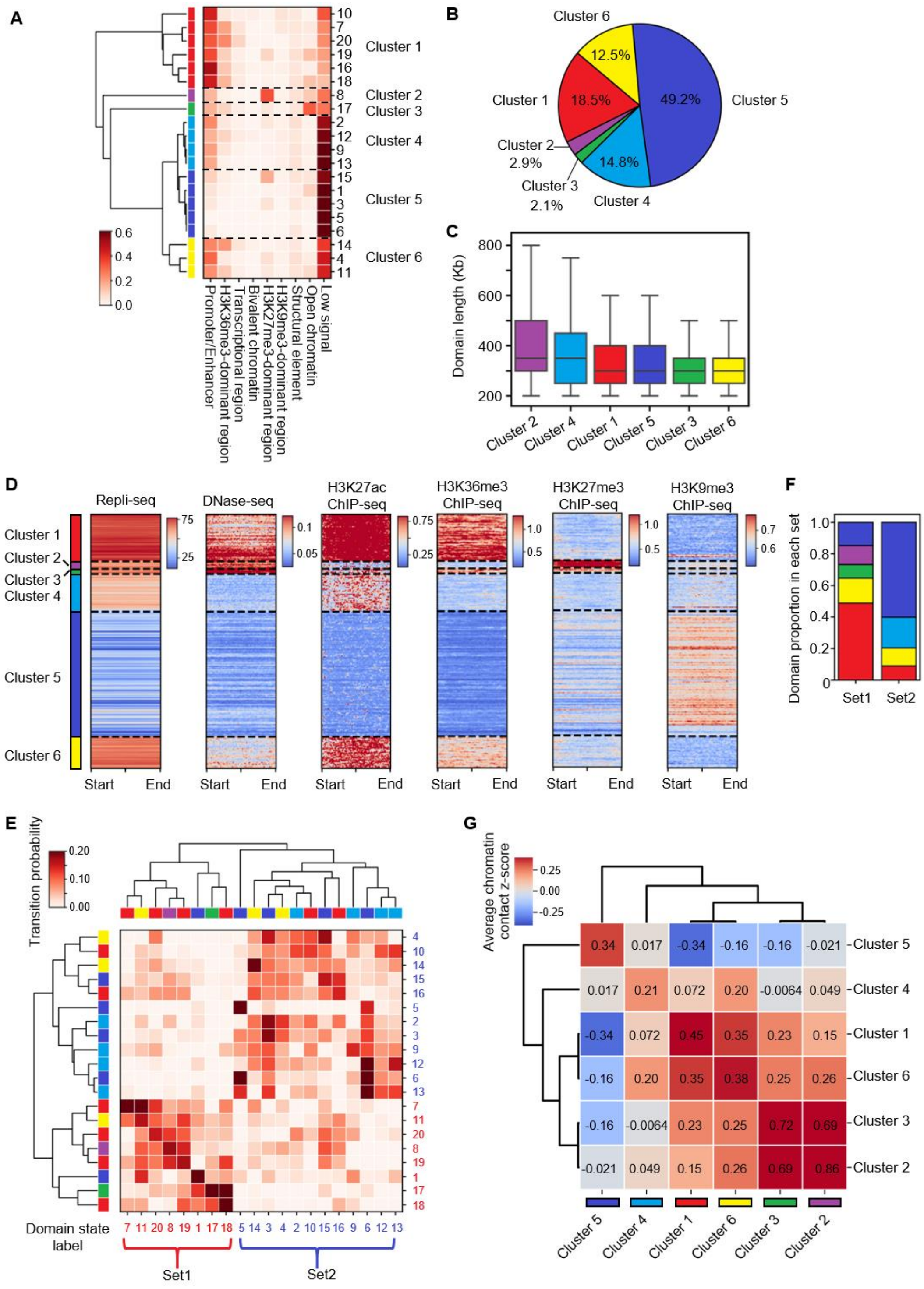


**Fig. S12. Define the types of domain clusters and analyze their biological characteristics. (A)** Hierarchical clustering of 20 domain states based on the proportions of nine types of regions within each type of domain. The 20 domain states are merged into six domain clusters. (**B**) The proportion of domains belonging to each of the six clusters among all topological domains. (**C**) The length distribution of topological domains within each of the six domain clusters. (**D**) The signal distribution of Repli-seq data, DNase-seq data, and ChIP-seq data of H3K27ac, H3K36me3, H3K27me3, and H3K9me3 within each domain in the six domain clusters. (**E**) The transition probability matrix between 20 domain states. Each row in the matrix represents the transition probability from the domain state corresponding to that row to other states corresponding to the columns. The color annotations for rows and columns indicate the domain cluster they belong to. The order of rows and columns has been rearranged based on hierarchical clustering with two clear domain-state sets. (**F**) The proportion of domains belonging to each domain cluster within two sets of domain states. (**G**) The average chromatin contact z-score between domains within the six domain clusters. A higher value means stronger chromatin contact between domains from the corresponding two clusters.

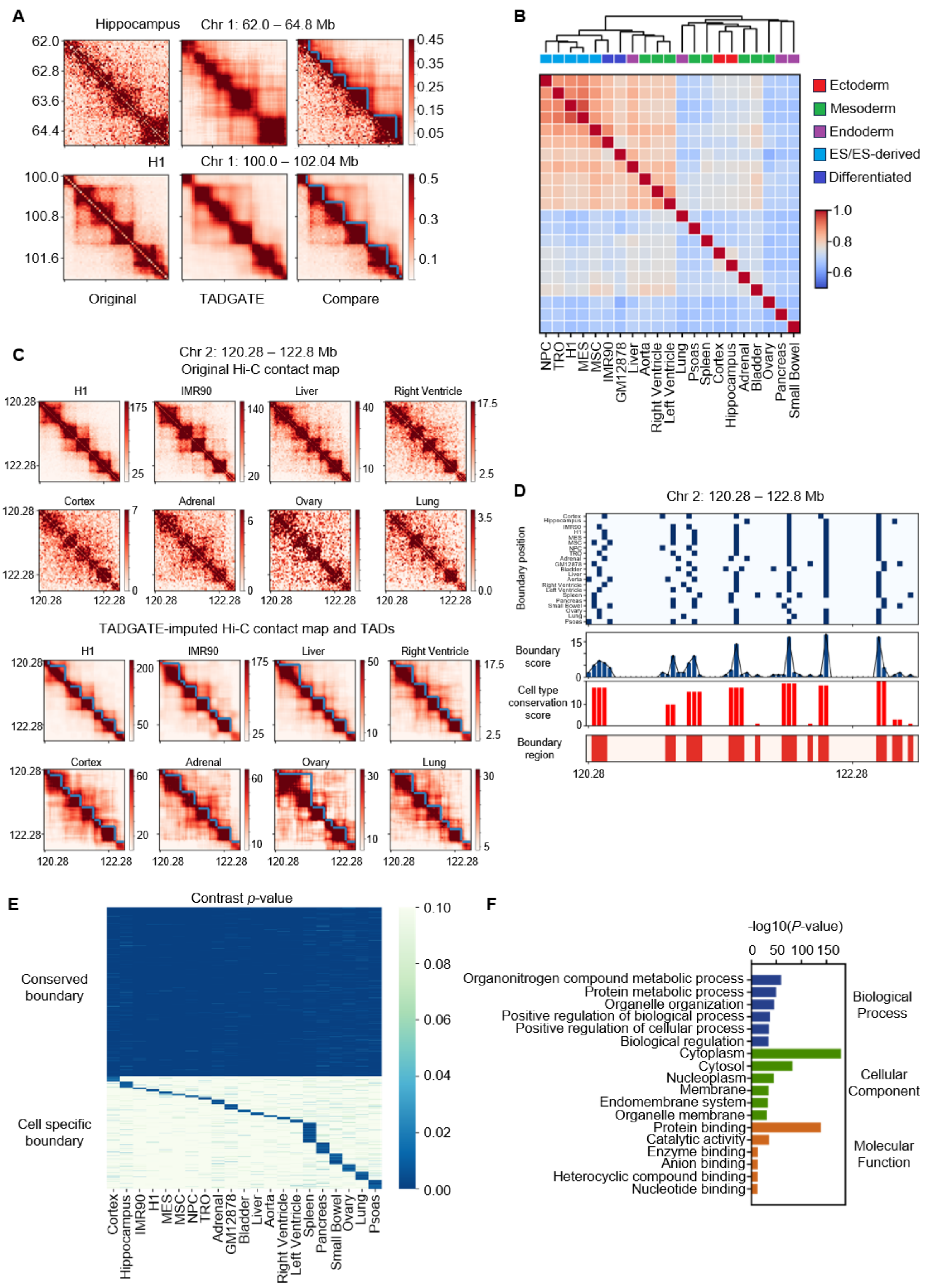


**Fig. S13**. **Analysis of Hi-C data of 21 human tissues and cell lines with TADGATE.** (**A**) Two example regions to show the original Hi-C contact maps of the hippocampus and Embryonic Stem Cell (H1), as well as the TADGATE-imputed contact maps and the corresponding TADs. In the third figure, the lower triangle represents the original map, while the upper triangle represents the TADGATE map. (**B**) Clustering results of all cell types based on the Spearman correlation coefficient of the Contrast Index with original contact maps for 21 tissues and cell lines. (**C and D**) A representative region to show the original Hi-C contact maps of 8 different cell types or tissues, the TADGATE-imputed maps, and the corresponding TADs (C). Based on the TADs identified in all cell types, the core boundary regions are defined, along with the conservation scores of these boundary regions across all cell types (D). (**E**) The contrast *p*-values of conserved boundaries for all 21 tissues or cell lines and specific boundaries for one cell type. (**F**) GO enrichment analysis for genes near the conserved boundaries. The top 6 GO terms in each category are shown.

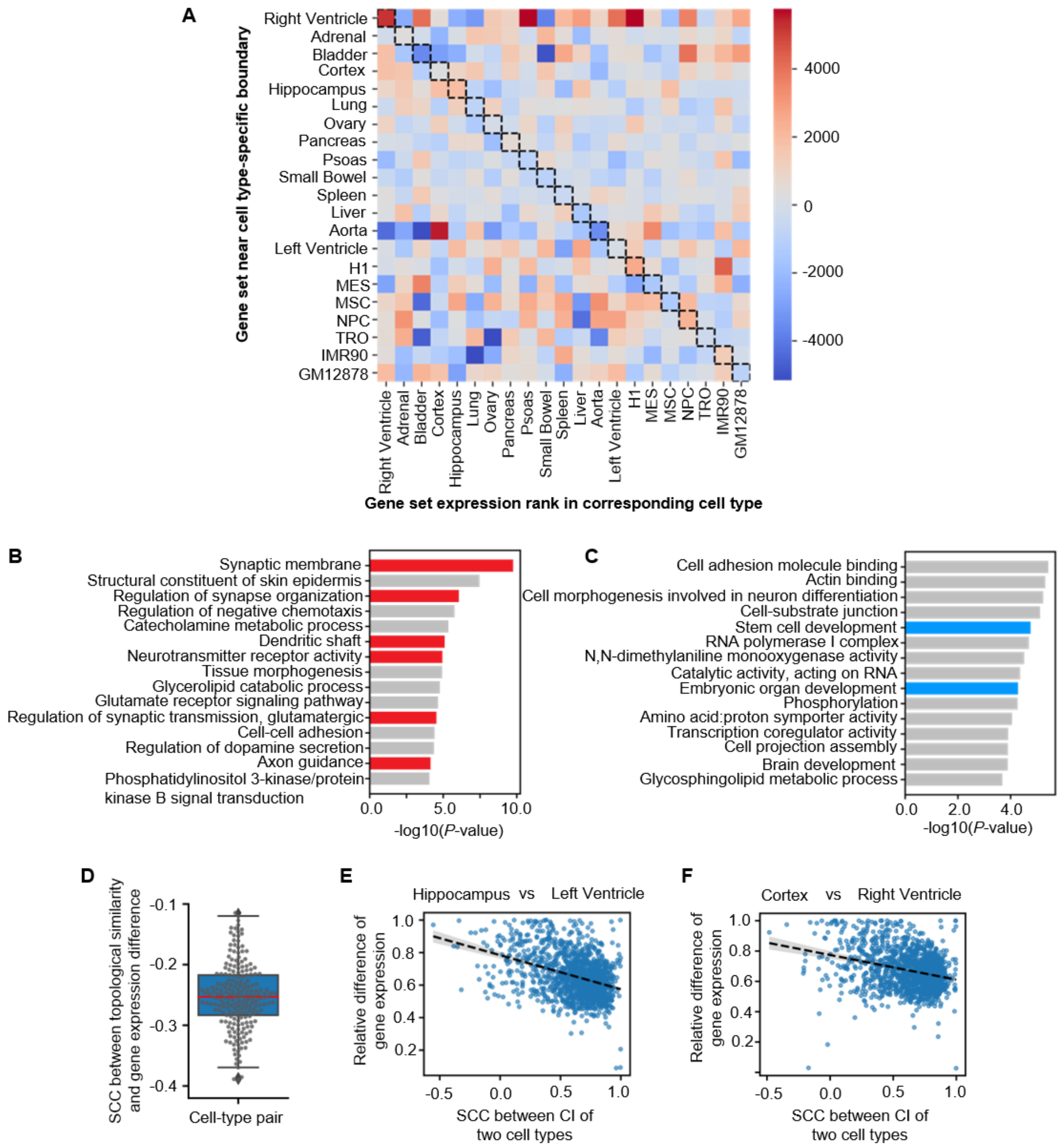


**Fig. S14**. **Analysis of genes around cell-type specific boundaries and the correlation between chromatin topological similarity and difference of gene expression.** (**A**) Rank analysis of gene expression levels for cell type-specific gene sets across different cell types. A higher rank indicates higher expression specificity of the gene set in the corresponding cell type. The diagonal values represent the rank of specific gene sets in their own cell type. (**B and C**) GO enrichment analysis for genes located in the nearby TADs delineated by brain-specific boundaries (B) or Embryo-specific boundaries (C). The GO terms related to the cell types are marked in red or blue. The top 15 GO terms are shown. (**D**) The Spearman correlation coefficient (SCC) between the contrast index (CI) similarity and gene expression difference for all cell-type pairs. (**E and F**) The Spearman Correlation Coefficient (SCC) between the contrast index (CI) profiles of two cell types and the relative gene expression difference between two cell types within each 2-Mb genomic interval. (E) for the Hippocampus and Left Ventricle and (F) for the Cortex and Right Ventricle. Each dot represents a 2-MB genomic interval and the linear regression is performed for SCC values and difference of gene expression.

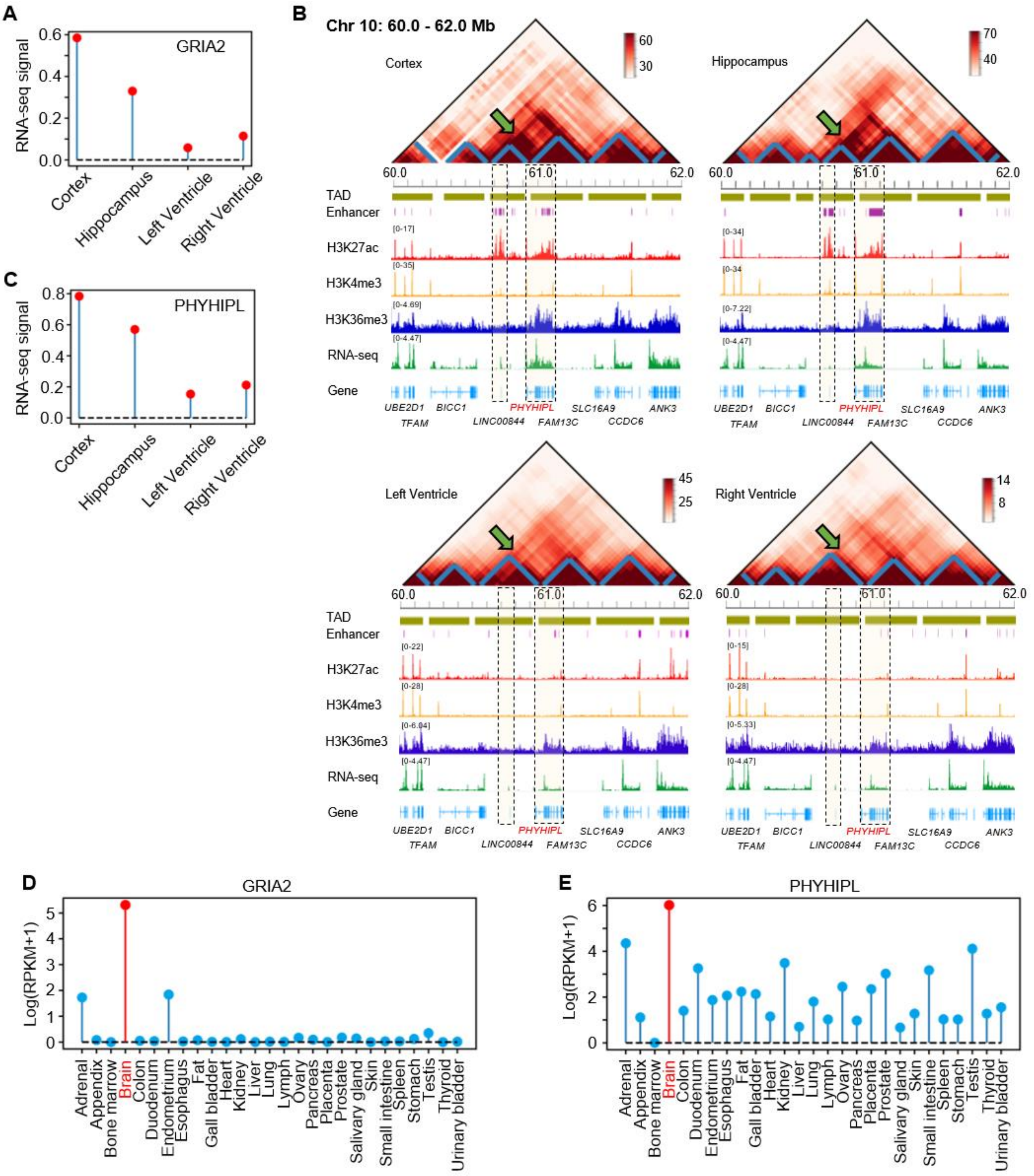


**Fig. S15**. **Analysis of the expression of genes *GRIA2* and *PHYHIPL* and the nearby structure of topological domains in different cell types.** (**A**) RNA-seq signal of gene *GRIA2* in the Cortex, Hippocampus, left and right Ventricle. (**B**) The comparison of TADGATE-imputed contact maps, chromatin topological domains, epigenetic signals, and RNA-seq signal around the gene *PHYHIPL* in the Cortex, Hippocampus, left and right Ventricle. (**C**) RNA-seq signal of gene *PHYHIPL* in the Cortex, Hippocampus, left and right Ventricle. (**D and E**) The expression of gene *GRIA2* or *PHYHIPL* across different human tissues.

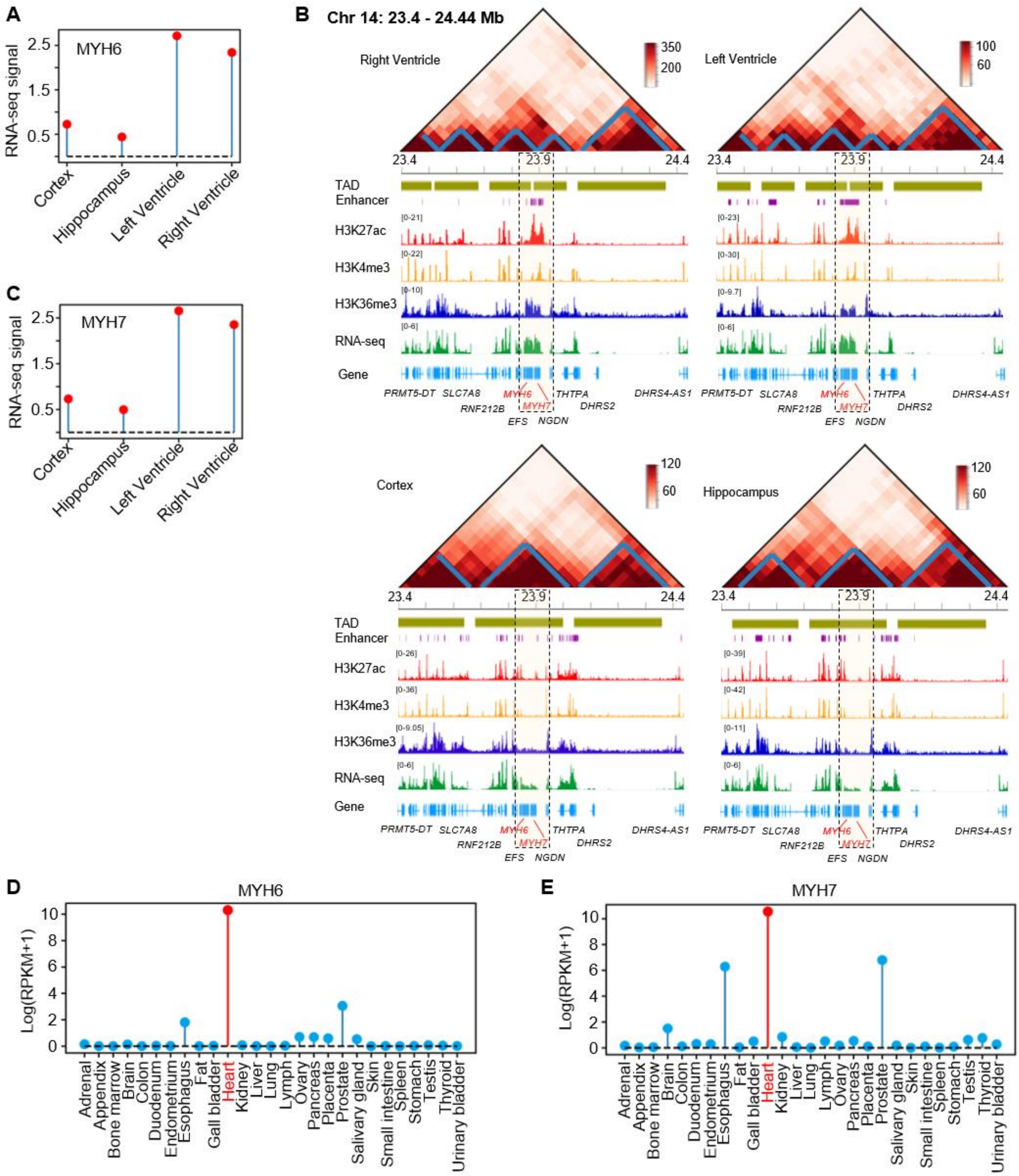


**Fig. S16**. **Analysis of the expression of genes *MYH6* and *MYH7* and the nearby structure of topological domains in different cell types.** (**A**) RNA-seq signal of gene *MYH6* in Cortex, Hippocampus, left and right Ventricle. (**B**) The comparison of TADGATE-imputed contact maps, chromatin topological domains, epigenetic signals, and RNA-seq signal around the genes *MYH6* and *MYH7* in the Cortex, Hippocampus, left and right Ventricle. (**C**) RNA-seq signal of gene *MYH7* in the Cortex, Hippocampus, left and right Ventricle. (**D and E**) The expression of gene *MYH6* or *MYH7* across different human tissues.
